# Acidity-induced dysfunction of CD8^+^ T cells is characterized by impaired IL-2 responsiveness and perturbations to mTORC1 signaling and c-Myc levels

**DOI:** 10.1101/2023.12.15.571931

**Authors:** Romain Vuillefroy de Silly, Laetitia Pericou, Bili Seijo, Isaac Crespo, George Coukos, Melita Irving

**Affiliations:** Ludwig Institute for Cancer Research, University of Lausanne and Department of Oncology, Lausanne University Hospital (CHUV), Lausanne, Switzerland

**Keywords:** Acidity, pH, CD8+ T cell, CTL, mTOR, c-Myc, IL-2, IL-2R signaling, Glutamine, Glutamate, Aspartate, Proline

## Abstract

CD8^+^ T cells play a critical role in cancer control but a range of barriers in the tumor microenvironment, including low pH, can impair their function. Here, we demonstrate that acidity dampens T-cell expansion mainly due to impaired IL-2 responsiveness, blunts cytokine secretion upon re-activation, and lowers the cytolytic capacity of CD8^+^ T cells expressing weak affinity TCR. We further reveal and dissect defects in both mTORC1 activity and c-Myc accumulation at low pH, the latter of which is largely due to proteasome-mediated degradation. In addition, lower intracellular levels of glutamine, glutamate and aspartate as well as elevated proline were noted, with no apparent impact on mTORC1 or c-Myc. Overall, low pH disrupts diverse intracellular signaling pathways as well as nutrient uptake/processing by T cells and we conclude that unless intracellular pH can be restored, multiple interventions will be required to overcome acidity-induced dysfunction.

## Introduction

Patient responses to cancer immunotherapies typically rely upon the re-invigoration of cytolytic T cells ^1, 2, 3, 4^ within a hostile tumor microenvironment (TME) ^5^. Along with powerful immune checkpoints such as the PD-1/PD-L1 axis and the presence of potently suppressive soluble factors (*e.g.* TGF-β, IL-10, adenosine and PGE-2) ^5, 6, 7^, the inherent physicochemical properties of the TME (*e.g.* stiffness, nutrient starvation, hypoxia, and acidity) represent major regulators of T-cell functions ^8, 9, 10, 11^. Extracellular acidification is a common property of tumors due to their reliance upon aerobic glycolysis (i.e., the Warburg effect) ^12, 13^ to meet their metabolic needs, leading to coupled efflux of lactate and protons, along with their tendency to over-express enzymes such as carbonic anhydrase ^14^. Interestingly, it has recently been shown that lymph nodes comprise acidic regions in which T cells themselves contribute to acidification of the extracellular milieu due to enhanced aerobic glycolysis upon activation^15^. It is well known that low pH can cause profound dysfunction of many cell types in the TME including effector CD8^+^ T cells ^11, 13, 15, 16, 17^, but the impact of acidity on cellular signaling and metabolism has not been comprehensively characterized. Elucidating mechanism(s) of action of low pH on T cells may contribute towards the development of more effective combinatorial immunotherapies and/or gene-engineering strategies ^18^ that can overcome this common barrier to T-cell activity in the TME. This notion is supported by the demonstration of improved tumor control in pre-clinical models upon modulation of acidity by administration of proton pump inhibitors ^19^ and bicarbonate ^20^.

Here, we have evaluated the impact of pH on effector CD8^+^ T-cell function upon re-activation *in vitro*, including of expansion/persistence as this is a major obstacle to most T-cell based immunotherapies ^1^. We identified defects in IL-2 responsiveness under acidic conditions and subsequently dissected how pH influences IL-2R signaling, the mTORC1 pathway, c-Myc activity, and metabolomic profiles. Overall, at low pH we reveal important changes to T-cell effector function and to IL-2 responsiveness, including intracellular signaling and transcription factor levels, and amino acid metabolism. Together, these findings implicate low pH in the TME as a major barrier to CD8^+^ T-cell responses and a critical target for improving responses to immunotherapy.

## Results

### Low pH impairs the function of effector CD8^+^ T cells upon re-activation

Within the TME anti-tumor CD8^+^ T cells are for the most part fully differentiated into an effector state (CTLs: cytotoxic T lymphocytes; ***Extended Data Fig. 1a***) ^1^. Hence, in order to explore the impact of low pH on T cells in the condition in which they would be found in tumors, we established an *in vitro* expansion protocol giving rise to effector murine T cells (***Extended Data Fig. 1c,d***). We chose pH7.4 as a control (i.e., the physiological pH) in comparison to pH7 and pH6.6. For OT-I TCR transgenic T cells which recognize the MHC class I/SIINFEKL peptide complex with high affinity, we observed no difference in cytolytic capacity at the different pH, even when modulating antigen density on tumor cells (***Fig. 1a,b***). In contrast, weaker affinity OT3 TCR CTLs demonstrated decreased cytolytic function at pH6.6 (***Fig. 1c***). We further evaluated cytokine production and proliferation, but, in order to prevent any CTL-extrinsic impact of acidity, the CTLs were re-activated with anti-CD3 antibodies instead of tumor cells. Upon re-stimulation at pH6.6 we observed a strong decrease in the production of all cytokines assessed (IFN-γ, IL-2 and TNF; ***Fig. 1d***). Interestingly, at low pH CTL proliferation was also dampened (***Fig. 1e***), independently of the stimulation strength (***Extended Data Fig. 2A***). We further observed that acidity prevented the increase in T-cell size and granularity normally induced upon activation at pH7.4 (***Extended Data Fig. 2b***).

**Fig. 1:**
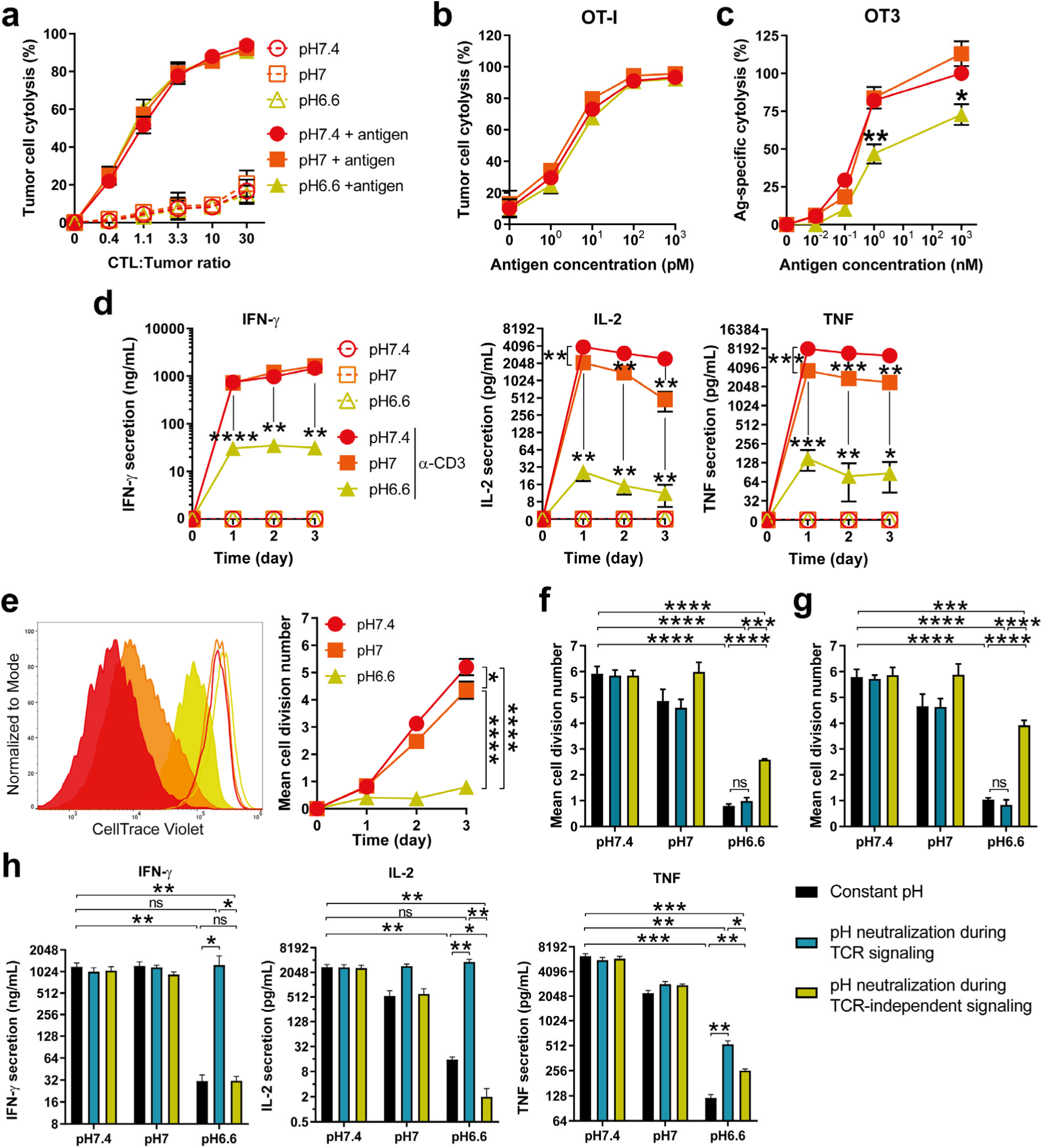
Low pH impairs the function of effector CD8+ T cells upon re-activation. **a,** Impact of pH on CTL killing capacities of OT-I as a function of CTL-to-tumor ratio. Varying numbers of OT-I CTLs were cocultured for 4.5 hours at three different pH (pH7.4, red circles, pH7, orange squares, or pH6.6, lime triangles) with C1498 tumor cells pulsed (solid lines and symbols), or not (dashed lines and empty symbols), with 1μM of antigen (minimal ovalbumin peptide epitope, SIINFEKL). Results show the mean percentage of tumor cell lysis ± SD of at least three (or two for 0.4 and 1.1 CTL:Tumor ratios) biological replicates from at least two independent experiments. **b,** Impact of pH on CTL killing capacities of OT-I as a function of antigen density. OT-I CTLs were cocultured for 4.5 hours at three different pH (pH7.4, red circles, pH7, orange squares, or pH6.6, lime triangles) with C1498 tumor cells pulsed with varying amounts of antigen at a 3.3 CTL-to-tumor ratio. Results show the mean percentage of tumor cell lysis ± SD of three biological replicates from two independent experiments. **c,** Impact of pH on CTL killing capacities of OT-3 as a function of antigen density. OT-3 CTLs were cocultured for 4.5 hours at three different pH (pH7.4, red circles, pH7, orange squares, or pH6.6, lime triangles) with C1498 tumor cells pulsed with varying amounts of antigen at a 3.3 CTL-to-tumor ratio. Results show the mean percentage of antigen-specific tumor cell lysis, normalized to the condition pH7.4 with 1μM of antigen, ± SEM of three biological replicates from two independent experiments. Statistic comparisons between pH6.6 and pH7.4: *p<0.05, **p<0.01 (Student’s paired *t*-test). **d,** Impact of pH on cytokine secretion profiles of CTLs. OT-I CTLs were cocultured for one, two, and three days in the presence (solid lines and symbols), or absence (dashed lines and empty symbols), of an agonistic anti-CD3 antibody (1μg/mL pre-coated plates) at different pH (pH7.4, red circles, pH7, orange squares, or pH6.6, lime triangles. Results show the mean secretion of IFN-γ, IL-2 and TNF-α ± SEM of at least three biological replicates from two independent experiments, as detected by ELISA from supernatants. *p<0.05, **p<0.01, ***p<0.001, ****p<0.0001 (Student’s paired *t*-test). **e,** Impact of pH on CTL proliferation upon anti-CD3 re-activation. OT-I CTLs were cultured for one, two, and three days in the presence of an agonistic anti-CD3 antibody (1μg/mL pre-coated plates) at different pH (pH7.4, red circles, pH7, orange squares, or pH6.6, lime triangles). Histograms show one representative experiment of CellTrace Violet dilution following re-activation with anti-CD3 (solid histograms), or without stimulation (empty histograms) at day 3. Bar graph shows the mean cell division number ± SEM of at least four biological replicates from at least two independent experiments. *p<0.05, ****p<0.0001 (Student’s paired *t*-test). **f,g,** Acidity affects CTL proliferation during TCR/CD3-independent signaling. **b,** OT-I CTLs were cultured for one day in anti-CD3 –coated plates (TCR/CD3 signaling step). The resulting cells and supernatants were transferred to an anti-CD3 –free plate and cultured for two further days (TCR/CD3-independent signaling step; cf. Supplementary Figure 2B). Cells were either: cultured during the TCR/CD3 signaling step and the TCR/CD3-independent signaling at constant pH (either pH7.4, pH7, pH6.6 – black bars, “Constant pH”), cultured at pH7.4 during the TCR/CD3 signaling step then at various pH during the TCR/CD3-independent signaling step (“pH neutralization during TCR signaling” – teal bars), or cultured at various pH during TCR/CD3 signaling step then at pH7.4 during the TCR/CD3-independent signaling step (“pH neutralization during TCR-independent signaling” –lime bars). Cell proliferation was measured by flow cytometry. (**g**) Same methodology as in (**f**), but adding exogenous murine IL-2 (200 IU/mL). Bar graphs show the mean cell division number +SEM of four biological replicates from two independent experiments. ns: not statistically significant, ***p<0.001, ****p<0.0001 (one-way repeated measures ANOVA, Turkey post-hoc test). **h,** Acidity influences cytokine secretion profile of CTLs during TCR signaling. The methodology was the same as in (**f**). Cytokine secretion was measured by ELISA. Bar graph shows the mean cytokine secretion of IFN-γ, IL-2 and TNF-α +SEM of four biological replicates from two independent experiments. ns: not statistically significant, *p<0.05, **p<0.01, ***p<0.001 (one-way repeated measures ANOVA, Turkey post-hoc test).

Next, we sought to evaluate at what stage proliferation is impaired upon re-activation of CTLs at low pH. CTL re-activation can be divided into two sequential steps: (i) a first TCR/CD3-dependent step leading to the production of cytokines, and, (ii) a second one that relies mainly upon autocrine IL-2 production and IL-2R signaling (***Extended Data Fig. 2c***). In order to determine the stage at which low pH is affecting CTL proliferation, we acidified/neutralized the pH at one step or the other (***Extended Data Fig. 2d***). We observed that pH neutralization during the IL-2R-dependent step improved CTL proliferation (***Fig. 1f***), even more so in the presence of exogenous IL-2 (***Fig. 1g***). Whereas, neutralizing pH during the TCR/CD3 signaling step restored both the cytokine secretion profile (***Fig. 1h***) and cell-size augmentation (***Extended Data Fig. 2e***). Taken together, our data show that acidity impacts both steps of CTL reactivation; low pH during TCR/CD3 triggering blunts cytokine secretion and cytolysis (the latter for low affinity TCR T cells), and acidic conditions at the IL-2R signaling phase impairs proliferation.

### Acidity abrogates IL-2-mediated proliferation of CTLs

To further validate that acidity dampens IL-2R signaling-driven proliferation we took advantage of the fact that CTLs maintain responsiveness to IL-2 without re-activation via TCR/CD3 triggering. We observed that even upon high dose IL-2 stimulation (200 IU/mL) acidity lowered CTL proliferation and viability (***Fig. 2a, Extended Data Fig. 3b***) as well as cell size and granularity (***Extended Data Fig. 3a,b***). Notably, pH6.6 did not preferentially block T cells in a particular phase of the cell cycle (***Extended Data Fig. 3c***).

**Fig. 2:**
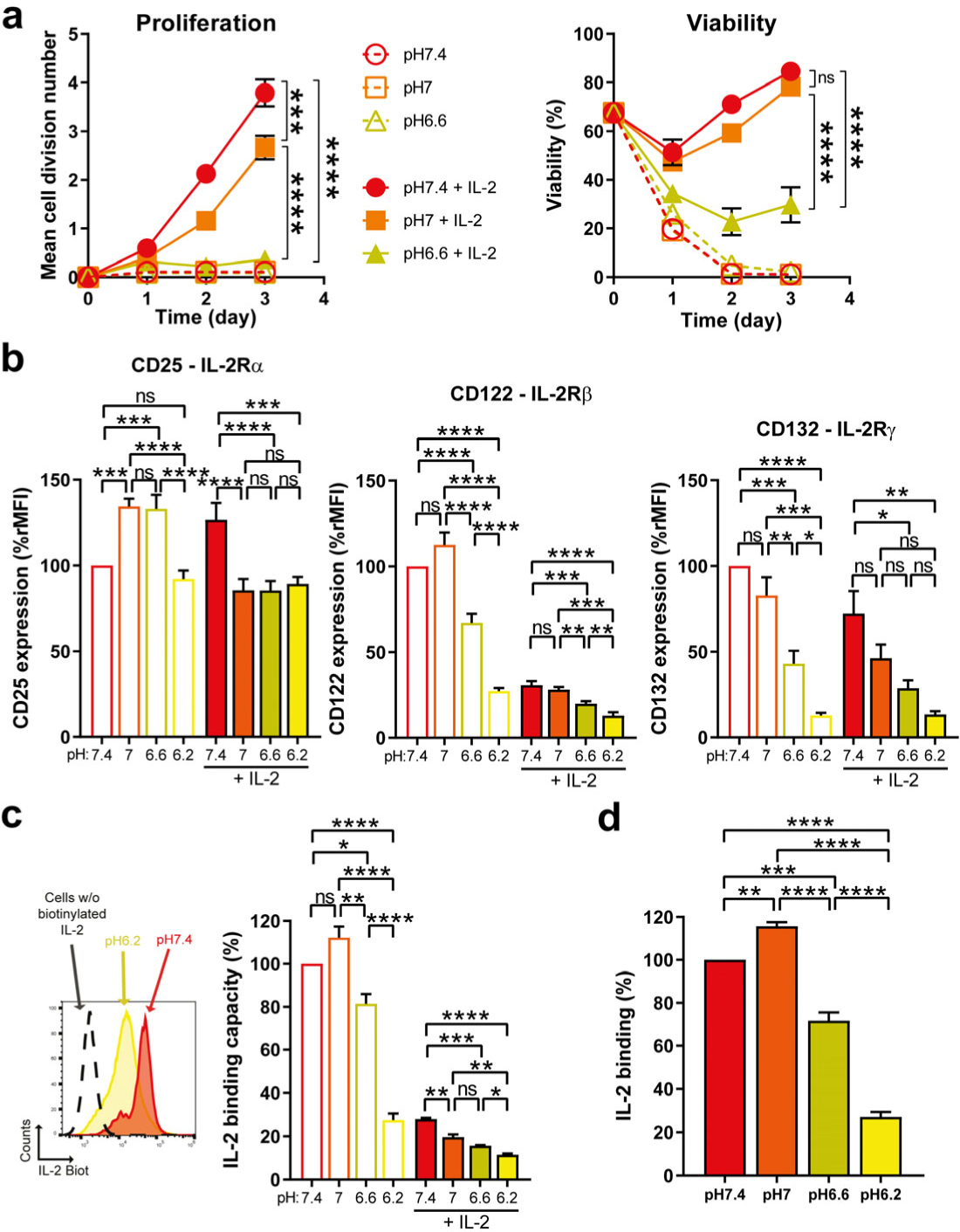
Acidity lowers IL-2-dependent proliferation of CTLs. **a,** Time-course of the pH impact proliferative and survival responses of CTLs to IL-2. OT-I CTLs were cultured for one, two, or three days at three different pH (pH7.4, red circles, pH7, orange squares, or pH6.6, lime triangles) in the presence (solid lines and symbols), or the absence (dashed lines and empty symbols), of exogenous murine IL-2 (200 IU/mL). Results show the mean cell division number, or viability, ± SEM of at least three biological replicates from at least two independent experiments. ns: not significant, ***p<0.001, ****p<0.0001 (Student’s paired *t*-test). **b,** Impact of pH on protein expression of IL-2R subunits. OT-I CTLs were cultured for one day at three different pH in the presence (solid bars), or absence (empty bars), of exogenous murine IL-2 (200 IU/mL). Results show the mean expression of the indicated subunits +SEM of three biological replicates from two independent experiments. Expression was calculated as ratio median fluorescence intensity and was normalized to the condition pH7.4 without exogenous IL-2. ns: not significant, *p<0.05, **p<0.01, ***p<0.001, ****p<0.0001 (one-way repeated measures ANOVA, Turkey post-hoc test). **c,** Impact of pH on the levels of IL-2R complexes that can bind IL-2. OT-I CTLs were cultured for one day at three different pH in the presence (solid bars), or absence (empty bars), of exogenous murine IL-2 (200 IU/mL). One representative experiment is shown as flow cytometry histograms: dashed line show CTLs cultured at pH7.4 without IL-2 for one day and not stained with biotinylated-IL-2, while red and yellow histograms represent CTLs cultured without IL-2 for one day at pH7.4 and pH6.2 and stained with biotinylated IL-2, respectively. Bar graph shows the mean IL-2 binding capacity +SEM of three biological replicates from two independent experiments. Binding capacity was calculated as ratio median fluorescence intensity of surface-detected biotinylated IL-2, normalized to the condition pH7.4 without exogenous IL-2. ns: not significant, *p<0.05, **p<0.01, ***p<0.001, ****p<0.0001 (one-way repeated measures ANOVA, Turkey post-hoc test). **d,** Impact of pH on the binding of IL-2 to IL-2R. OT-I CTLs stained with biotinylated IL-2 for 30 minutes at 4°C in PBS at various pH. Bar graph shows the mean IL-2 binding +SEM of three biological replicates from two independent experiments. Binding was calculated as ratio median fluorescence intensity of surface-detected biotinylated IL-2, normalized to the condition pH7.4. **p<0.01, ***p<0.001, ****p<0.0001 (one-way repeated measures ANOVA, Turkey post-hoc test).

We subsequently sought to evaluate potential changes to the cell-surface expression of the IL-2R complex at low pH. Briefly, the IL-2R comprises an α (CD25), β (CD122), and γ (CD132) chain and can engage IL-2 by a βγ (intermediate affinity) or αβγ (high affinity) complex. The latter can be found on activated T cells upon upregulation of the α-chain. For both complexes, IL-2R signaling is mediated by the cytoplasmic domains of the IL-2Rβ and IL-2Rγ chains ^21, 22, 23^. After 24 hours incubation at pH6.6 we observed a decrease in the β and γ subunits at the cell-surface, and this was even more pronounced at pH6.2 (***Fig. 2b***). Notably, we also detected lower levels of the β- and γ-chains when the cells were pre-incubated with IL-2, presumably due to steric hindrance to antibody-binding and/or internalization of the chains (***Fig. 2b***). To evaluate if there was also a decrease in functional IL-2R complexes at low pH, we incubated the cells with biotinylated IL-2 (bio-IL-2) to allow detection of its binding (***Fig. 2c***). For this assay, competition by previously bound non-bio-IL-2 probably precludes detection of all cell-surface IL-2R complexes. Interestingly, we observed a decrease in bio-IL-2/IL-2R binding capacity at pH6.6, which was even more pronounced at pH6.2 (***Fig. 2d***). Overall, since only the most extreme acidity tested (i.e., pH6.2) was associated with decreased IL-2R levels and highly impaired IL-2/IL-2R binding, we conclude that the profound decrease in proliferation and viability of T cells cultured at pH6.6 is not due to lowered IL-2/IL-2R engagement.

Finally, in order to exclude that the impact of acidity on T cells resulted from indirect effects such as precipitation/inactivation of components in the medium, we analyzed T-cell proliferation with medium previously acidified and then re-adjusted to pH7.4 with NaOH. No proliferation defect was observed for T cells upon restoration of physiologic pH. In addition, because the experimental set-up involves HCl to reach the desired pH, we further tested if an increase in osmolarity could impact T-cell proliferation (by addition of NaCl) but this was not the case (***Extended Data Fig. 3d***).

### Acidity blunts IL-2R signaling, and specifically lowers mTORC1 activity and c-Myc levels

Given that IL-2/IL2-R complex formation is only modestly reduced at pH 6.6 but IL-2-mediated proliferation is nonetheless significantly impaired, we next sought to evaluate changes to intracellular signaling components. Briefly, IL-2/IL-2R complex formation first triggers the phosphorylation of Janus Kinases, JAK1 and JAK3. The JAKs then phosphorylate tyrosine residues in the intracellular domains of the IL-2R to generate docking sites for the signal transducer and activator of transcription (STAT) factors (e.g., STAT5) which dimerize and translocate to the nucleus and induce gene expression programs. The JAKs also initiate PI3K/Akt/mTORC1 and MAPK/ERK signaling pathways, and promote c-Myc transcriptional activity (***Fig. 3a***) ^22^. While we observed an important pH-dependent reduction in the phosphorylation of JAK1, JAK3 and STAT5 (***Fig. 3b***) at pH6.2, the decrease was only modest at pH6.6. However, although upstream transducers were slightly disturbed at pH6.6, we observed a strong impact on both the mTORC1 pathway (as evidenced by phosphorylation of p70S6K, S6 and 4E-BP1) ^24, 25^ and c-Myc levels (***Fig. 3c***). In contrast, the MAPK/ERK pathway was not impaired and even increased at later time-point, presumably due to cell-cycle differences (***Fig. 3c***).

**Fig. 3:**
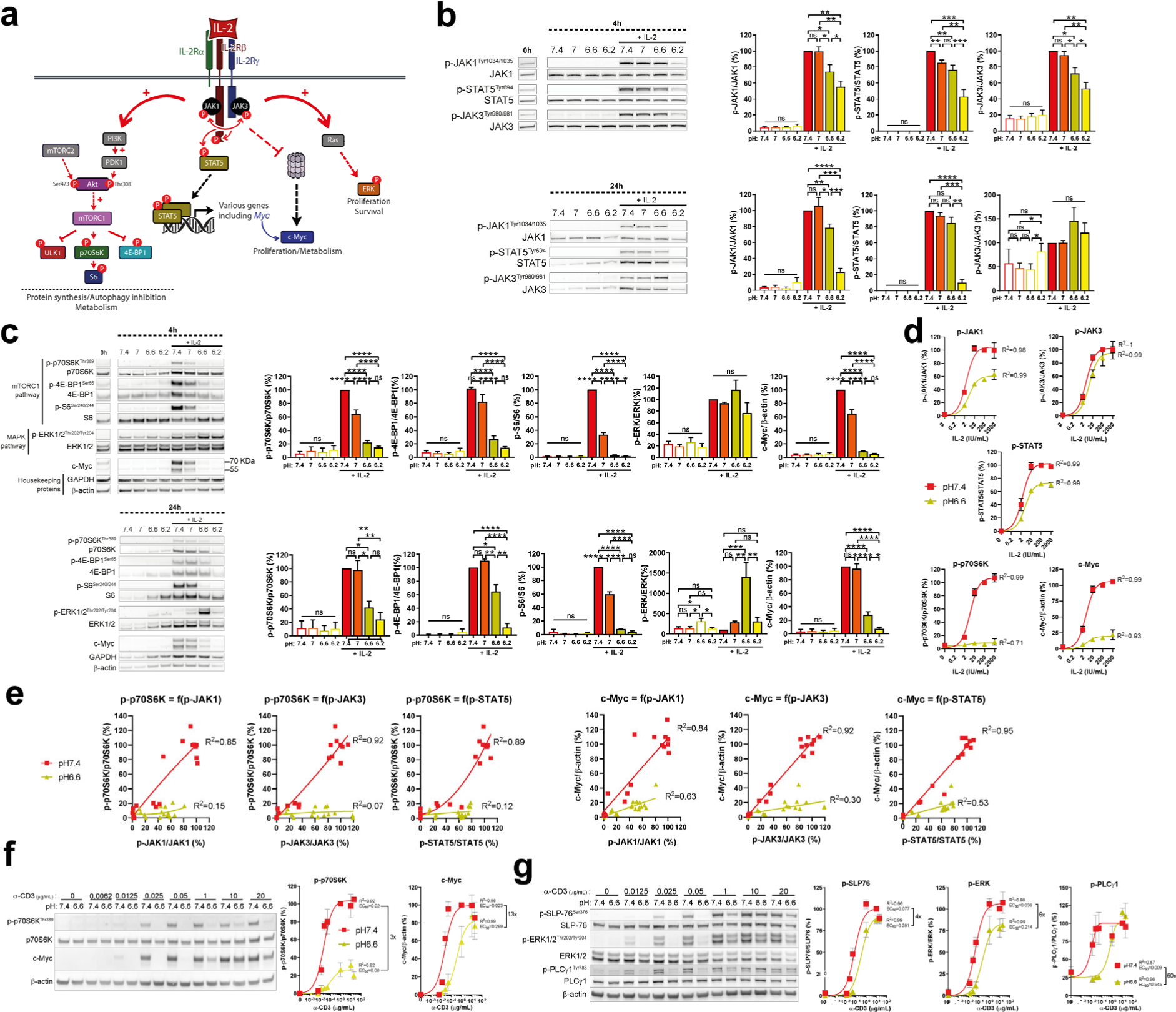
Low pH affects IL-2R signaling and activation threshold, and specifically mTORC1 and c-Myc levels. **a,** Simplified scheme of the IL-2R signaling. **b,** IL-2R signaling is disturbed at lower pH. OT-I CTLs were cultured at various pH in the presence, or the absence of exogenous murine IL-2 (200 IU/mL) for 4 or 24 hours (without prior starving). One representative western blot is shown. Bar graphs show the mean phosphorylation status relative to the total protein of interest normalized to the condition “pH7.4 + IL-2” +SEM of at least five biological replicates from at least two independent experiments. ns: not significant, *p<0.05, **p<0.01, ***p<0.001, ****p<0.0001 (one-way repeated measures ANOVA, Turkey post-hoc test). **c,** Low pH disturbs IL-2 –induced mTORC1 pathway and c-Myc levels. OT-I CTLs were cultured at various pH in the presence, or the absence of exogenous murine IL-2 (200 IU/mL) for 4 or 24 hours (without prior starving). One representative western blot is shown. Bar graphs show the mean phosphorylation status, or total levels, of the indicated molecule relative to the total protein of interest, or to β-actin, normalized to the condition “pH7.4 + IL-2” ± SEM of at least five biological replicates from at least two independent experiments. ns: not significant, *p<0.05, **p<0.01, ***p<0.001, ****p<0.0001 (two-way ANOVA, Turkey post-hoc test). **d,** IL-2R signaling dose-response. OT-I CTLs were cultured at various pH (pH7.4: red squares; pH6.6: lime triangles) in the presence, or the absence of various exogenous doses of murine IL-2 for 4 hours. Results show the mean phosphorylation status, or total levels, of the indicated molecule relative to the total protein of interest, or to β-actin, normalized to the condition “pH7.4 + IL-2 200 IU/mL” ± SEM of four biological replicates from two independent experiments. Corresponding correlation curves together with associated R^2^ are displayed. **e,** Correlation between the activation of first signaling transducers and mTORC1 pathway target activation, or c-Myc levels. Results show individual values obtained from the experiments displayed in (**d**). Corresponding correlation curves together with associated R^2^ are shown. **f,** mTORC1 activity and c-Myc levels upon TCR/CD3 signaling dose-response at low pH. OT-I CTLs were cultured at various pH (pH7.4: red squares; pH6.6: lime triangles) in the presence of various doses of coated-anti-CD3 antibodies for 4 hours. One representative western blot experiment is shown. Results show the mean phosphorylation status of p70S6K, or c-Myc total levels, normalized to the condition “pH7.4 + anti-CD3 1μg/mL” ± SEM of three biological replicates from two independent experiments. Corresponding correlation curves together with associated R^2^ and EC50 are displayed. **g,** TCR/CD3 signaling dose-response at low pH. OT-I CTLs were cultured at various pH (pH7.4: red squares; pH6.6: lime triangles) in the presence of various doses of coated-anti-CD3 antibodies for 4 hours. One representative western blot experiment is shown. Results show the mean phosphorylation status of SLP-76, ERK or PLCγ1 normalized to the condition “pH7.4 + anti-CD3 10μg/mL” ± SEM of three biological replicates from two independent experiments. Corresponding correlation curves together with associated R^2^ and EC50 are displayed.

We subsequently assessed the activation of various pathways that could lead to mTORC1 inhibition, including (i) stress response (p38, eiF2α), (ii) energy homeostasis (AMPK), (iii) Wnt/Hedgehog signaling (GSK-3β), (iv) amino acid starvation (eiF2α) and, (v) cAMP response (CREB/CREM) ^26, 27, 28^, but we did not observe any notable differences at low pH (***Extended Data Fig. 4a,b***). In line with the described role of mTORC1 as a major regulator of protein synthesis, we observed that acidic pH was associated with lower protein content in the cells (***Extended Data Fig. 4c***). We further preformed RNA sequencing on CTLs cultured for 24 hours at pH7.4 versus pH6.6 in the presence of IL-2, and gene set enrichment analysis identified c-Myc targets and the mTORC1 pathway as downregulated at low pH (***Extended Data Fig. 4d***).

Next, we evaluated the impact of low pH on IL-2R signaling and mTORC1/c-Myc at 4 hours, a time point at which cell viability and apoptosis are not impacted by acidic conditions (***Extended Data Fig. 5a***). We sought to elucidate whether the modest decrease in IL-2/IL-2R binding and JAK/STAT phosphorylation at pH6.6 could account for the significant decreases in phosphorylation of molecules downstream of mTORC1, and to lower c-Myc levels. We performed an IL-2 dose-response experiment at pH7.4 versus pH6.6 and observed impaired mTORC1 activation and lower c-Myc at levels seemingly disproportionate to a global decrease in IL-2R signaling (***Fig. 3d***). Indeed, by plotting individual values of mTORC1 pathway activation (p-p70S6K as a surrogate) and c-Myc levels as a function of JAK1/JAK3/STAT5 phosphorylation (surrogates of IL-2R signaling) as correlation curves (***Fig. 3e***), we observed no overlap for OT-I TCR T cells cultured at pH6.6 versus pH7.4. These data suggest that the decrease in mTORC1 and c-Myc levels are not due to lower IL-2R signaling. The same was observed for wild-type polyclonal C57BL/6 CTLs (***Extended Data Fig. 5b,c***).

Recently, Gaggero et. al., reported the development of a human IL-2 switch mutant able to preferentially engage IL-2R at low pH, thereby restoring signaling and proliferation ^29^. We generated this mutant in-house but could not reproduce their findings for either murine (***Extended Data Fig. 6a,b***) nor human CD8^+^ T cells (***Extended Data Fig. 6c,d***). The IL-2 switch mutant is described to specifically rescue binding to the IL-2Rα subunit at low pH. However, we observed that although knockout of IL-2Rα lowered all signaling parameters analyzed upon IL-2 stimulation, acidity still had a negative impact (***Extended Data Fig. 6e-h***).

Finally, we performed a dose-response curve for anti-CD3 antibody stimulation to determine if mTORC1 and c-Myc levels are also altered at low pH during the first step of CTL re-activation (***Extended Data Fig. 1a***). Both mTORC1 and c-Myc levels were blunted at low pH, but at high levels of anti-CD3 antibody c-Myc levels could be restored (***Fig. 3f***). We thus investigated TCR/CD3 signaling pathway molecules SLP-76, ERK and PLCγ1 (***Fig. 3g***) and found that at low pH the EC_50_ to reach maximum phosphorylation levels (as achieved at pH7.4) increased. This indicates that low pH augments the activation threshold, resulting in the c-Myc response observed.

### Low pH leads to decreased *Myc* transcription and to proteasome-mediated c-Myc degradation

We next questioned whether mTORC1 and c-Myc cross-regulate one another and thus performed a kinetic analysis. We observed a drop in mTORC1 as early as 5 to 15 minutes post low-pH exposure (even in the absence of exogenous IL-2) while c-Myc only decreased after an hour in an IL-2 dependent manner (***Fig. 4a***). Although mTORC1 is a major regulator of protein synthesis (*i.e.* its inhibition at low pH could account for the decrease in c-Myc protein levels), its inhibition by rapamycin was associated with only a minimal decrease in c-Myc (***Fig. 4b***) thereby indicating that there is no interplay between mTORC1 and c-Myc under acidic conditions, at least during the first four hours. We further explored *Myc* mRNA levels and found them decreased at pH6.6 (***Fig. 4c***), at least in part due to lower p-JAK1/JAK3/STAT5 signaling (***Fig. 4d***). The lack of correlation between c-Myc protein and Myc mRNA levels (***Fig. 4e***) indicated a potential role for post-transcriptional regulation of c-Myc levels ^30^, a mechanism oftentimes dependent upon its phosphorylation status at T58 (GSK-3β-dependent; p58 promotes proteasomal degradation) and S62 (ERK-dependent; pS62 improves stability). We did not observe a disturbance in pT58/pS62 upon acidic treatment, and GSK-3β inhibition (CHIR99021/GSKi) did not upregulate c-Myc levels, but we found that proteasome inhibition (MG-132) restored c-Myc levels (***Fig. 4f***). Taken together, our results indicate that drop in c-Myc levels under acidic conditions is mainly caused by increased proteasomal degradation, and is also due to lower *Myc* transcription as a result of weaker IL-2R signaling.

**Fig. 4:**
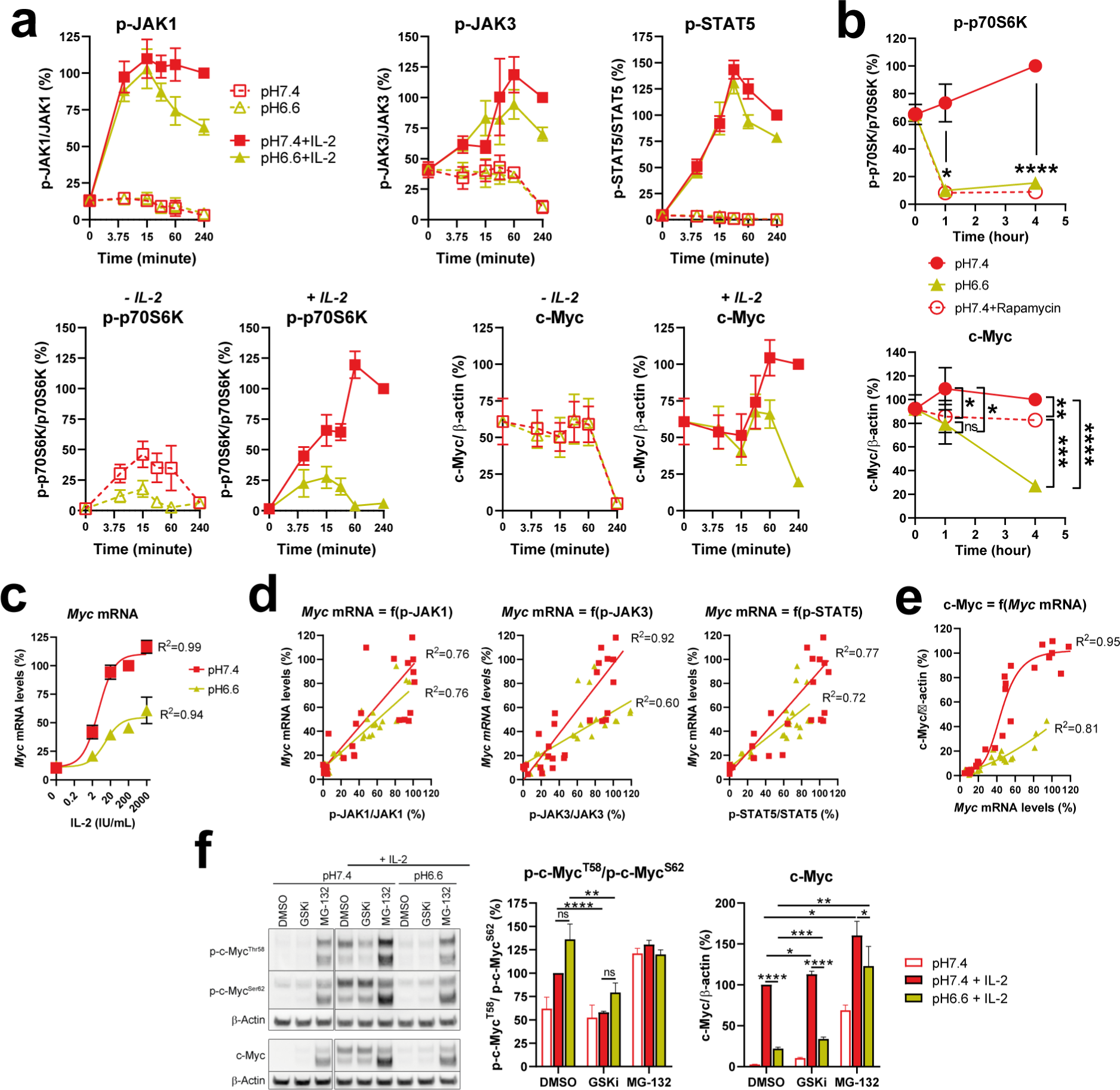
Low pH leads to decreased *Myc* transcription and to proteasome-mediated c-Myc degradation. **a,** Time-course of IL-2R signaling. OT-I CTLs were cultured at various pH in the presence (pH7.4: red squares, solid lines; pH6.6: lime triangles, solid lines), or the absence (pH7.4: empty red squares, dashed lines; pH6.6: empty lime triangles, dashed lines) of exogenous murine IL-2 (200 IU/mL) for various time. Results show the mean ± SEM of four biological replicates from two independent experiments. **b,** Impact of mTOR inhibition of c-Myc levels. OT-I CTLs were cultured at various pH with exogenous murine IL-2 (200 IU/mL) in the absence (pH7.4: red circles, solid lines; pH6.6: lime triangles, solid lines), or in the presence (pH7.4: empty red circles, dashed lines) of rapamycin 10nM for various time. Results show the mean ± SEM of four biological replicates from two independent experiments. **c,** *Myc* mRNA transcription. OT-I CTLs were cultured for 4 hours with various concentrations of exogenous murine IL-2 at pH7.4 (red squares and line) or at pH6.6 (lime squares and line). Line graph displays the mean percentage of *Myc* mRNA levels relative to *Actb* as a function of extracellular IL-2, normalized to the condition “pH7.4 + IL-2 200 IU/mL” ± SEM of four biological replicates from two independent experiments. A correlation curve and R^2^ for each pH is displayed. **d,** *Myc* mRNA levels as a function of IL-2R signaling. OT-I CTLs were cultured for 4 hours with various concentrations of exogenous murine IL-2 at pH7.4 (red squares and line) or at pH6.6 (lime squares and line). Dot plots display the individual percentage of *Myc* mRNA levels relative to *Actb* as a function of extracellular JAK1, JAK3 or STAT5 phosphorylation, normalized to the condition “pH7.4 + IL-2 200 IU/mL” from four biological replicates from two independent experiments. A correlation curve and R^2^ for each pH is displayed. **e,** c-Myc as a function of *Myc* mRNA levels. OT-I CTLs were cultured for 4 hours with various concentrations of exogenous murine IL-2 at pH7.4 (red squares and line) or at pH6.6 (lime squares and line). Dot plots display the individual percentage of c-Myc protein as a function of *Myc* mRNA levels, normalized to the condition “pH7.4 + IL-2 200 IU/mL” from four biological replicates from two independent experiments. A correlation curve and R^2^ for each pH is displayed. **f,** Acidity does not modify c-Myc phosphorylation but decreases c-Myc levels via the proteasome. OT-I CTLs were cultured for 4 hours with or without exogenous murine IL-2 (200 IU/mL) at pH7.4 or pH6.6 with, or without, the proteasome inhibitor MG-132 (10μM) or the GSK inhibitor CHIR99021 (2μM - GSKi). A representative western blot is shown. Bar graph on the left hand displays the mean ratio p-c-MycT58/p-c-MycS62 normalized to the condition “pH7.4 + IL-2 + DMSO” +SEM of four biological replicates from two independent experiments. Bar graph on the right hand shows the total levels of c-Myc relative to β-actin, normalized to the condition “pH7.4 + IL-2 + DMSO” +SEM of four biological replicates from two independent experiments. ns: not significant, *p<0.05, **p<0.01, ***p<0.001, ****p<0.0001 significant (Student’s paired *t*-test).

### TSC2 knockout augments mTORC1 activity at low pH but does not restore CTL proliferation

We next sought to explore mechanisms limiting mTORC1 activity at low pH. Interestingly, although mTORC1 inhibition with rapamycin recapitulated proliferative defects observed under acidic conditions (***Fig. 5a***), the phosphorylation status of Akt (an important activator of mTORC1; ***Fig. 5b***), was not altered at pH6.6 (***Fig. 5c***). Moreover, although Akt inhibition lowered mTORC1 activation and c-Myc accumulation, we did not detect a decrease in the phosphorylation status of direct Akt targets GSK-3β and FoxO, while PRAS40 phosphorylation was only modestly lowered. Subsequently, by CRISPR/Cas9 we individually knocked out PRAS40 as well as TSC2, the latter of which is a major negative regulator of mTORC1 that is also controlled by Akt (***Fig. 5d***). PRAS40 was not found to play a role in the regulation of mTORC1 activity, but loss of TSC2 resulted in mTORC1 activation, even in the absence of IL-2 and regardless of pH. The addition of IL-2 further increased mTORC1 activity in TSC2 KO CTLs at pH7.4 in an Akt-independent manner (***Fig. 5e***), but had little impact on mTORC1 at pH6.6. Despite the elevated mTORC1 activity at low pH in TSC2 KO cells, there was not an increase in CTL proliferation (***Fig. 5f***).

**Fig. 5:**
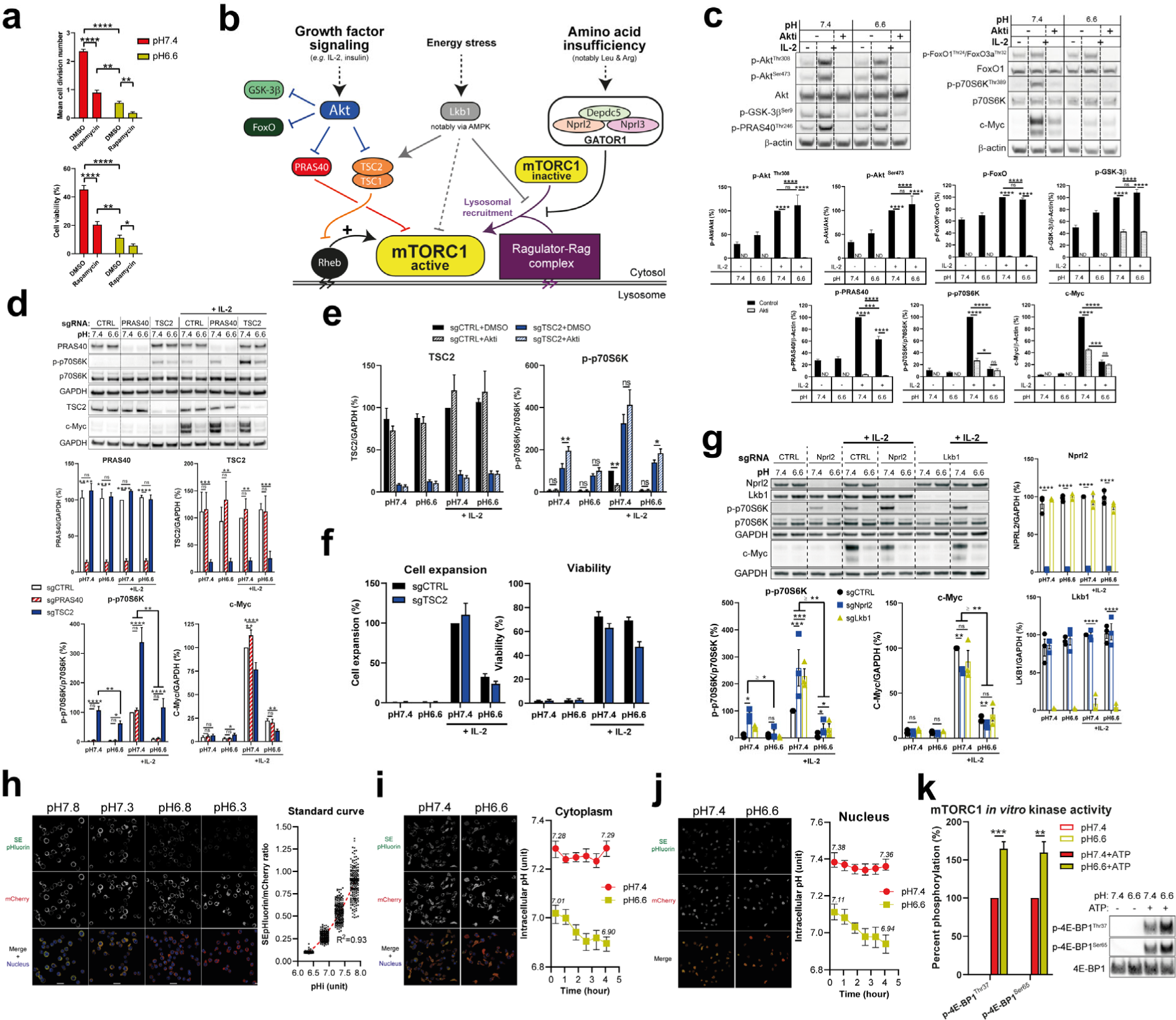
Knocking out TSC2 improves mTORC1 activity at low pH, while acidity lowers intracellular pH. **a,** mTORC1 inhibition leads to proliferation and viability defects following IL-2 stimulation. OT-I CTLs were cultured for three days in the presence (“Rapamycin”), or absence (“DMSO”), of rapamycin 10nM at pH7.4 (red bars) or pH6.6 (lime bars) with exogenous murine IL-2 (200 IU/mL). Results show the mean cell division number, or viability, +SEM of three biological replicates from two independent experiments. *p<0.05, **p<0.01, ****p<0.0001 (one-way repeated measures ANOVA, Turkey post-hoc test). **b,** Simplified scheme of mTORC1 activity regulation. mTORC1 is recruited to the lysosome via the Ragulator-Rag complex where it can eventually be activated by Rheb. Growth factor signaling leads to Akt activation that phosphorylates PRAS40 and TSC2 allowing to block their capacity to inhibit mTORC1 activity. Amino acid sufficiency prevents GATOR1 complex from impeding mTORC1 recruitment to lysosomes. Energy stress leads to Lkb1 activation that can lower mTORC1 activity via several mechanisms (mostly involving AMPK) including TSC2 activation, Ragulator-Rag inhibition and mTORC1 inhibiting phosphorylation. **c,** Acidity does not inhibit phosphorylation of direct Akt targets. OT-I CTLs were cultured for 4 hours at pH7.4 or pH6.6 with, or without, exogenous murine IL-2 (200 IU/mL), in the presence, or absence (“Control”, DMSO), of Akt1/2 inhibitor (10μM-Akti). One representative western blot is shown. Bar graphs show the mean phosphorylation status, or total levels, of the indicated molecule relative to the total protein of interest, or to β-actin, normalized to the condition “pH7.4 + IL-2” +SEM of four biological replicates from two independent experiments. ns: not significant, *p<0.05, ***p<0.001, ****p<0.0001 (two-way ANOVA, Turkey post-hoc test); ND: not determined. **d,** TSC2 knockout improves mTORC1 activity. OT-I x CRISPR/Cas9 CTLs were transduced with retroviruses encoding a negative control, a PRAS40 or a TSC2 sgRNA, and were cultured for 4 hours at pH7.4 or pH6.6 with, or without, exogenous murine IL-2 (200 IU/mL). One representative western blot from two membranes of the same samples is shown. Bar graphs show the mean phosphorylation status, or total levels, of the indicated molecule relative to the total protein of interest, or to GAPDH, normalized to the condition “pH7.4 + IL-2 sgCTRL” +SEM of four biological replicates from two independent experiments. ns: not significant, *p<0.05, **p<0.01, ***p<0.001, ****p<0.0001 (two-way ANOVA, Turkey post-hoc test). **e,** TSC2 knockout improves mTORC1 activity. OT-I x CRISPR/Cas9 CTLs were transduced with retroviruses encoding a negative control, a PRAS40 or a TSC2 sgRNA, and were cultured for 4 hours at pH7.4 or pH6.6 with, or without, exogenous murine IL-2 (200 IU/mL) in the presence, or absence (“DMSO”), of Akt1/2 inhibitor (10μM-Akti). Bar graphs show the mean phosphorylation status, or total levels, of the indicated molecule relative to the total protein of interest, or to GAPDH, normalized to the condition “pH7.4 + IL-2 sgCTRL DMSO” +SEM of four biological replicates from two independent experiments. ns: not significant, *p<0.05, **p<0.01 (Student’s paired *t*-test). **f,** TSC2 knockout does not improve CTL proliferation. OT-I x CRISPR/Cas9 CTLs were transduced with retroviruses encoding a negative control (black bars) or a TSC2 (blue bars) sgRNA, and were cultured 4 days at pH7.4 or pH6.6 with, or without, exogenous murine IL-2 (200 IU/mL). Results show the relative expansion (normalized to the condition “pH7.4 + IL-2 sgCTRL”), or viability, +SEM of at least four biological replicates from at least two independent experiments. **g,** Nprl2 and Lkb1 knockouts do not improve mTORC1 activity at low pH. OT-I x CRISPR/Cas9 CTLs were transduced with retroviruses encoding a negative control, a Nprl2 or a Lkb1 sgRNA, and were cultured for 4 hours at pH7.4 or pH6.6 with, or without, exogenous murine IL-2 (200 IU/mL). One representative western blot from two membranes of the same samples is shown. Bar graphs show the mean phosphorylation status, or total levels, of the indicated molecule relative to the total protein of interest, or to GAPDH, normalized to the condition “pH7.4 + IL-2 sgCTRL” +SEM of three biological replicates (except for conditions pH7.4 and pH6.6 sgLkb1 without IL-2: two) from two independent experiments. ns: not significant, *p<0.05, **p<0.01, ***p<0.001, ****p<0.0001 (two-way ANOVA, Turkey post-hoc test). **h,** Standard curve to detect intracellular pH. OT-I CTLs were engineered to express SEpHluorin-mCherry. CTLs were cultured in high K^+^ buffer containing Valinomycin/Nigericin with defined pH, allowing to obtain equilibrium between extracellular and intracellular pH. Dot plot graph shows the ratio of fluorescence of SEpHluorin/mCherry per cell excluding nucleus at each pH (>100 cell/pH) acquired by live confocal microscopy. A correlation curve was obtained using the median fluorescence for each pH and was used as a standard curve. Representative images for each pH are displayed. **i,** Extracellular acidity lowers intracytoplasmic pH. OT-I CTLs were engineered to express SEpHluorin-mCherry. CTLs were cultured at pH7.4 (red circles) or pH6.6 (lime squares) in the presence of exogenous murine IL-2 (200 IU/mL) for various time, and were acquired by live confocal microscopy. Line graph shows the median ratio of fluorescence of SEpHluorin/mCherry per cell excluding nucleus ± 95% CI of at least 150 cell per time-point per pH from one out of two independent experiments. Representative images for each pH are displayed. **j,** Extracellular acidity lowers nuclear pH. OT-I CTLs were engineered to express SEpHluorin-mCherry. CTLs were cultured at pH7.4 (red circles) or pH6.6 (lime squares) in the presence of exogenous murine IL-2 (200 IU/mL) for various time, and were acquired by live confocal microscopy. Line graph shows the median ratio of fluorescence of SEpHluorin/mCherry per cell excluding extra-nucleus signal ± 95% CI of at least 150 cell per time-point per pH from one out of two independent experiments. Representative images for each pH are displayed. **k,** mTORC1 kinase activity *in vitro* is not lowered at pH6.6. Kinase activity of recombinant mTORC1 complexes (mTOR/RAPTOR/MLST8) at pH7.4 or pH6.6 was assessed by determining phosphorylation status of recombinant 4E-BP1 upon a 10 minutes reaction at 30°C in the presence, or absence, of ATP. One representative Western blot is shown. Results show the mean phosphorylation status of 4E-BP1 normalized to the condition “pH7.4 + ATP” +SEM of four independent experiments. **p<0.01, ***p<0.001 (Student’s t-test).

Rheb is directly inhibited by TSC2, and Rheb overexpression has been demonstrated to facilitate unhindered mTORC1 encounter and activation ^31, 32^. We overexpressed Rheb and observed similar effects as for TSC2 KO cells, albeit less pronounced (***Extended Data Fig. 7a***). In addition to growth factors and Akt, mTORC1 activity is regulated by amino acid insufficiency and energy stress via GATOR1 and the Lkb1/AMPK axis, respectively (***Fig. 5b***). Interestingly, KO of Nprl2 which blocks the GATOR1 axis led to activation of mTORC1 even in the absence of IL-2, but not at pH6.6 (***Fig. 5g***). Also, despite increasing mTORC1 activity upon addition of IL-2 at pH7.4, Nprl2 and Lkb1 KOs minimally augmented mTORC1 activity under acidic conditions (***Fig. 5g***).

### Extracellular acidity promotes intracellular acidification

Having demonstrated thus far that acidic conditions (pH6.6) blunt T-cell function including IL-2 mediated proliferation with apparent marginal impairment to IL-2/IL-2R binding and signaling but strong independent reduction in mTORC1 activity and c-Myc levels, we reasoned that there must be important intracellular acidification disruptive to multiple independent cellular processes. We sought an experimental approach for which it would be possible to distinguish acidification within the cytoplasm versus nucleus, a large organelle in CTLs. Briefly, we engineered T cells to express SEpHluorin/mCherry ^33^, a fusion protein comprising SEpHluorin (a pH-sensitive mutant of GFP) and mCherry (pH-insensitive) and allowing one to infer intracytoplasmic and nuclear pH in parallel by ratiometric measurements via confocal microscopy by comparison with a standard curve (***Fig. 5h***). Interestingly, we observed that intracellular pH was rapidly (≤ 20min) acidified (***Fig. 5i***) down to ∼pH7.0 (vs ∼ pH7.3 in the control group, corresponding to a H^+^ increase of more than 80%) upon pH6.6 treatment, and that it dropped to ∼pH6.90 (corresponding to a H^+^ increase of more than 140%) at the end of the experiment (4 hours). Acidic conditions also lowered nuclear pH to the same extent and with the same kinetics (***Fig. 5j***) despite that the nucleus had a slightly more alkaline pH at baseline (+0.1 pH unit). We speculated that low pH may directly impair the enzymatic activity of mTORC1. However, we found that the ability of mTORC1 complexes to phosphorylate purified 4E-BP1 was in fact higher in vitro at pH6.6 than at pH7.4 (***Fig. 5k***). Indeed, mTORC1 functions in close proximity to lysosomes which are highly acidic (pH4.5-5) ^34^ which may explain our observation.

### Acidity lowers intracellular glutamine/glutamate/aspartate levels and promotes proline accumulation

Finally, we sought to explore potential differences in the metabolome of T cells under acidic conditions which could potentially play a role in cellular function ^35^. Both mTORC1 and c-Myc are known regulators of amino acid homeostasis and can themselves be controlled by amino acid levels ^36, 37^. Interestingly, under acidic conditions we observed a consistent decrease in the intracellular levels of glutamine and glutamate, and an increase in proline content (***Fig. 6a,b***). Since mTORC1 activation and c-Myc levels have been previously described to be modulated by glutamine ^37, 38^, we assessed whether low glutamine levels could be causing the effects observed at low pH. We confirmed that glutamine deprivation from the medium mimics impaired CTL proliferation and mTORC1/c-Myc patterns observed under acidic conditions (***Fig. 6c,d***). However, a glutamine dose-response experiment revealed that mTORC1/c-Myc patterns at low pH were matched at 1000-fold less (2μM) glutamine in the culture medium (***Fig. 6e***) while merely a 10-fold decrease (200μM) in extracellular glutamine completely abrogated intracellular glutamine detection in CTLs (***Fig. 6f***). Plotting mTORC1 and c-Myc as a function of intracellular glutamine or glutamate (***Extended Data Fig. 7b***) further indicate that low intracellular glutamine/glutamate under acidic conditions does not dictate mTORC1/c-Myc patterns but rather could be a consequence of them. We observed that increasing doses of extracellular glutamine lowered intracellular levels of serine and threonine (***Extended Data Fig. 7c***), which might reflect CTL reliance on ASCT2, a transporter which imports glutamine while exporting serine and threonine ^39, 40^.

**Fig. 6:**
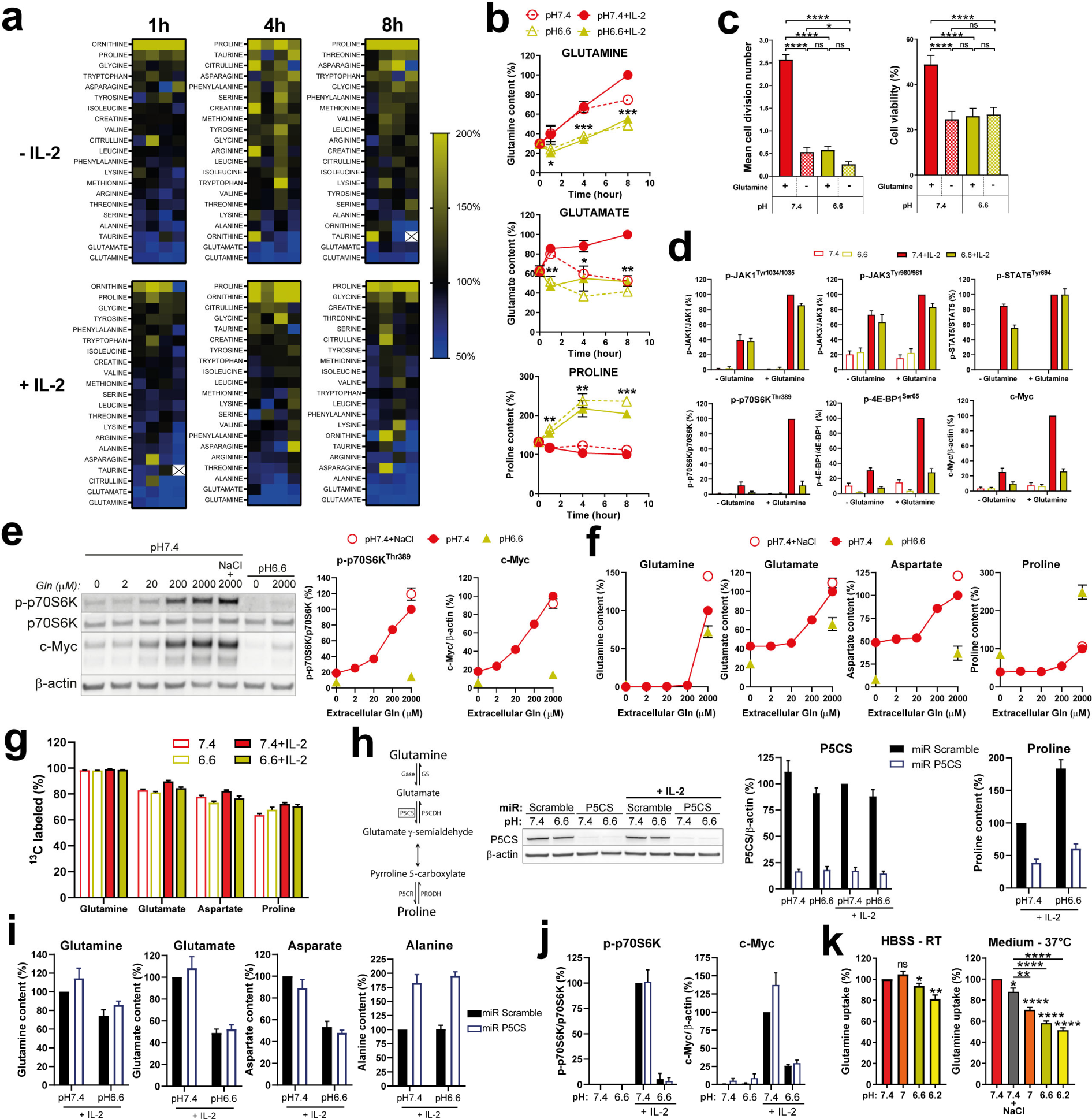
Acidity lowers intracellular glutamine/glutamate/aspartate levels and promotes proline accumulation. **a,** Heat map of intracellular amino acid contents as a function of pH, time, and IL-2 stimulation. Intracellular levels of amino acids were quantified from OT-I CTLs that were cultured for various time in the presence, or absence, of exogenous murine IL-2 (200 IU/mL) at pH7.4 or pH6.6. Results show the percent level of amino acid as compared to the matched pH7.4 condition using a color code, as outlined on the ladder. Each square show one biological replicate, with a total of four from two independent experiments. White square with a cross inside means the amino acid was not detected. **b,** Time-course of intracellular levels of glutamine, glutamate, and proline. Intracellular levels were quantified as in (**a**). OT-I CTLs were cultured for various time in the presence (solid symbols and lines), or absence (empty symbols and dashed lines), of exogenous murine IL-2 (200 IU/mL) at pH7.4 (red circles) or pH6.6 (lime triangles). Results show the mean intracellular level of the amino acid of interest (normalized to the condition “pH7.4 + IL-2 8h”) ± SEM of four biological replicates from two independent experiments. ns: not significant, *p<0.05, **p<0.01, ***p<0.001 (Student’s paired *t*-test). **c,** Glutamine deprivation inhibits CTL proliferation and viability. OT-I CTLs were cultured for three days with exogenous murine IL-2 (200 IU/mL) at pH7.4 or pH6.6 in the presence, or absence, of glutamine (under the glutamax form, 4mM). Results show the mean cell division, or cell viability, +SEM of at least four biological replicates from at least two independent experiments. ns: not significant, *p<0.05, ****p<0.0001 (two-way ANOVA, Turkey post-hoc test). **d,** Glutamine deprivation lowers mTOR activation and c-Myc accumulation. OT-I CTLs were cultured for 4 hours with (solid bars), or without (empty bars), exogenous murine IL-2 (200 IU/mL) at pH7.4 (red bars) or pH6.6 (lime bars) in the presence, or absence, of glutamine (2mM). Results show the mean phosphorylation status, or total levels, of the indicated molecule relative to the total protein of interest, or to β-actin, normalized to the condition “pH7.4 + IL-2” ± SEM of four biological replicates from two independent experiments. **e,** Dose-dependent impact of glutamine on phosphorylation of p70S6K and c-Myc levels. OT-I CTLs were cultured for 4 hours with exogenous murine IL-2 (200 IU/mL) at pH7.4 or pH6.6 in the presence of various quantities of glutamine, and in presence or absence of NaCl 31mM. One representative western blot out of four biological replicates from two independent experiments is shown. Line graphs display the mean relative phosphorylation status of p70S6K, c-Myc levels ± SEM of the pooled data, as a function of extracellular glutamine content. **f,** Dose-dependent impact of exogenous glutamine on amino acid levels. OT-I CTLs were cultured for four hours with exogenous murine IL-2 (200 IU/mL) at pH7.4 or pH6.6 in the presence of various quantities of glutamine, and in presence or absence of NaCl 31mM. Results show the mean intracellular amino acid content normalized to the condition “pH7.4 + 2000μM Gln” ± SEM of four biological replicates (the same replicates as the one used for (**e**)) from two independent experiments. **g,** Intracellular glutamine, glutamate, aspartate and proline are coming from extracellular glutamine. OT-I CTLs were cultured for 4 hours with isotopic ^13^C-glutamine (2mM) in the presence (solid bars), or absence (empty bars), of exogenous murine IL-2 (200 IU/mL) at pH7.4 (red bars) or pH6.6 (lime bars). Results show the mean proportion of the indicated intracellular amino acid that incorporated ^13^C +SEM of four biological replicates from two independent experiments. **h,** Knockdown of P5CS prevents proline accumulation. OT-I CTLs transduced with a control miR (Scramble) or a P5CS-targeting miR were cultured for 4 hours with, or without, exogenous murine IL-2 (200 IU/mL) at pH7.4 or pH6.6. One representative western blot displaying P5CS protein levels is shown. Bar graph on the left hand displays the mean ratio p-c-MycT58/p-c-MycS62 normalized to the condition “pH7.4 + IL-2 + DMSO” +SEM of four biological replicates from two independent experiments. Bar graph on the left hand displays the mean P5CS protein levels normalized to the condition “pH7.4 + IL-2 miR Scramble” +SEM of four biological replicates from two independent experiments. Bar graph on the right hand shows proline levels normalized to the condition “pH7.4 + IL-2 miR Scramble” +SEM of four biological replicates from two independent experiments. **i,** P5CS knockdown does not improve glutamine/glutamate/aspartate levels at low pH, but increase alanine accumulation. Intracellular levels of amino acids were determined from the same CTLs as in (**h**). Results show the mean amino acid level normalized to the condition “pH7.4 + IL-2 miR Scramble” +SEM of four biological replicates from two independent experiments. **j,** P5CS knockdown does not improve mTORC1 activity and c-Myc levels at low pH. p70S6K phosphorylation and c-Myc levels were determined from the same CTLs as in (**h**). Results show the mean level normalized to the condition “pH7.4 + IL-2 miR Scramble” +SEM of four biological replicates from two independent experiments. **k,** Acidity lowers glutamine uptake in CTLs. OT-I CTLs were cultured for 20 minutes either in HBSS at room temperature, or in CTL medium with IL-2 (200 IU/mL) at 37°C. Amino acid uptake assay was carried out by adding tritiated glutamine at 10μCi/mL (equivalent to 165nM). Results show the mean glutamine uptake normalized to the condition “pH7.4” +SEM of eight biological replicates from at least three independent experiments. ns: not significant, *p<0.05, **p<0.01, ****p<0.0001 (Student’s paired t-test).

While glutamine is considered the major source of glutamate ^41^, proline and aspartate can arise from glutamine/glutamate conversion ^42^. Thus, we next performed tracer experiments with ^13^C isotopic glutamine. Of note, within four hours upwards of 50% of some TCA cycle metabolites (*i.e.* citrate, fumarate, and malate) were generated from extracellular glutamine (***Extended Data Fig. 7d***). Importantly, we confirmed that more than 80% of the glutamate pool, more than 70% of the aspartate pool, and at least 65% of the proline pool were derived from glutamine (***Fig. 6g***). We questioned if the decrease in intracellular levels of glutamine/glutamate/aspartate could be due to increased conversion to proline. While a miR-based knockdown (of more than 80%) of P5CS, a rate-limiting enzyme involved in proline conversion (***Fig. 6h***) blocked an increase in proline levels at low pH, there was no rescue of glutamine/glutamate/aspartate levels (***Fig. 6i***), mTORC1/c-Myc patterns (***Fig. 6j***), or cell proliferation and viability (***Extended Data Fig. 6e***). Upon P5CS knockdown there was, however, an increase in alanine accumulation (***Fig. 6i***). Finally, utilizing tritiated glutamine we determined that its uptake was pH-dependent (***Fig. 6j***). Notably, the assay had to be performed using complete medium at 37°C (instead of HBSS at room temperature) ^43^ in order to see a clear impact, thus suggesting that acidity disturbs amino acid uptake and/or export (including of glutamine) via competition for transporters. All perturbations to CTLs at low pH are summarized in a graphical abstract in ***Extended Data Fig. 8*.**

## Discussion

Here, we have comprehensively explored the impact of acidity, a common suppressive feature of solid tumors, on effector CD8^+^ T cells. It has been previously shown that the cytolytic capacity of CTLs is blunted at low pH ^44, 45^ and we confirmed this to be the case for T cells expressing weak but not high affinity TCRs. In line with previous studies, we also observed reduced cytokine secretion (IFN-γ, IL-2 and TNF) and proliferation upon CTL re-activation under acidic conditions. Several mechanisms have been proposed to explain how acidity impairs CTLs including CD3ζ downregulation, lower CD25 (IL-2Rα) levels, upregulation of CTLA-4 and PD-1, lower phosphorylation of Akt, ERK, STAT5, p38 and JNK, and/or activation of proton-sensing receptors (*e.g.* ASICs, TDGA8, OGR1, TRPV) ^19, 20, 45, 46, 47, 48^, but previous studies are awash with contradictory findings.

To gain deeper mechanistic insight into acidity-induced T-cell dysfunction, we considered CTL re-activation in two stages, a first TCR/CD3-dependent step during which cytokines are generated, and a second one that relies mainly upon autocrine IL-2 production and IL-2R signaling. We found that although acidity inhibited IL-2 production during CTL re-activation, impaired proliferation was mostly due to diminished IL-2 responsiveness in a TCR/CD3-independent manner. We observed that levels of the IL-2R subunits (mostly β and γ), functional IL-2R complex, IL-2 binding capacity, and the phosphorylation of upstream signaling molecules (i.e., JAK1, JAK3 and STAT5) dropped under acidic conditions. However, at a moderate acidity of pH6.6, IL-2R signaling was only marginally impacted, and the MAPK/ERK pathway not at all. In contrast, at pH6.6 we observed that both mTORC1 activation (as evaluated by phosphorylation of p70S6K, S6, 4E-BP1 and ULK1) and c-Myc accumulation were reduced, independently of each other and of a global IL-2R signaling decrease.

mTOR is a central hub integrating growth factor signaling and nutrient availability to regulate critical cellular processes including protein and nucleotide synthesis, and ultimately T-cell fate ^25^. Interestingly, although the mTORC1 inhibitor rapamycin ^23, 49^ blunted CTL proliferation (i.e., suggesting that impaired proliferation at low pH is associated with decreased mTORC1 activity), we observed that Akt activation (a kinase which can activate mTORC1) and its downstream targets including GSK-3β, FoxO1 and PRAS40 were not disrupted under acidic conditions. GATOR1 (involved in amino acid sensing) and Lkb1/AMPK pathways were similarly not involved in the reduced mTORC1 activation at low pH. Moreover, we observed no phosphorylation or accumulation of the transcription factors CREB and CREM, suggesting that proton-sensing receptors which often signal through the cAMP/PKA pathway and can lead to mTORC1 inhibition ^50, 51^ are not involved in reduced mTORC1 activity under acidic conditions. Similarly, Wu et al. recently ruled out participation of several proton-sensing receptors in acidity-mediated suppression of T cells ^15^.

mTORC1 inhibition at low pH has been described in fibroblasts and tumor cells ^52, 53, 54^ and it has been proposed that acidification within the cytoplasm causes lysosome dispersion thereby preventing mTORC1 co-localization with its activator Rheb ^54^. We observed that CTL culture at low pH causes rapid acidification within both the cytoplasmic and nuclear compartments, but given the small size of the cytoplasm in T cells we could not demonstrate such a mechanism (data not shown). However, the fact that TSC2 (a Rheb inhibitor) knockout promoted mTORC1 activity argues against lysosome dispersion taking place in CTLs. While TSC2 knockout enforced mTORC1 activity at low pH, this was not sufficient to restore CTL proliferation upon IL-2 stimulation. Finally, we questioned if low pH could directly inhibit mTORC1, but the kinase activity of the purified complex in vitro was not disrupted at pH6.6. Hence, we conclude that along with its well-known roles in sensing nutrient and energy sufficiency, mTORC1 acts as a sensor of extracellular pH, an important parameter for integration in order to adapt cellular physiology to the microenvironment.

The transcription factor c-Myc plays critical roles during T-cell activation and IL-2-mediated proliferation, it regulates metabolic programs including glutaminolysis, and its inactivation prevents the proliferation of T cells ^55, 56, 57^. Under acidic conditions we observed a slight decrease in IL-2/IL-2R binding and in JAK/STAT activation, and consequently a modest decrease in *Myc* transcription. Interestingly, although we did not observe changes in phosphorylation of T58 or S62, the residues mostly commonly involved in driving proteasomal degradation or stabilization of c-Myc, respectively,^30^ we found that c-Myc accumulation could be restored upon proteasome inhibition.

As observed during IL-2R signaling, acidity lowered mTORC1 activity and c-Myc levels upon re-activation of CTLs. Interestingly, in contrast to mTORC1, we noticed c-Myc decrease was related to increased activation threshold imposed by acidity and could be compensated by increasing the activation stimulus…Given the multifunctional roles of both c-Myc and mTORC1 including in protein synthesis (which is needed for the generation of cytokines, granzymes and perforin, etc…), it is likely that impaired CTL effector function at low pH is at least in part due to their perturbation. Further work is warranted to delineate the precise mechanisms by which c-Myc and mTORC1 are inhibited under acidic conditions.

Finally, we questioned if low pH could cause changes in the metabolism of amino acids. Indeed, amino acids play a crucial role in T-cell function and they are tightly connected to mTORC1 and c-Myc activities. Under acidic conditions we observed a drop in intracellular levels of glutamine, glutamate and aspartate, whereas proline accumulated. Glutamine is a nonessential amino acid and its uptake and catabolism are highly induced in active T cells to provide intermediate molecules for different pathways of biosynthesis as well as substrates for the mitochondria ^58^. For example, during glutaminolysis its carbon backbone can be converted to α-ketoglutarate to maintain homeostasis of the tricarboxylic-acid cycle (TCA), or to lactate that generates NAD and NADPH ^59^. Notably, insufficient glutamine can inhibit T-cell proliferation, growth and cytokine production ^60^.

Using isotopic glutamine, we determined that glutamate, aspartate and proline in CTLs are mostly derived from extracellular glutamine. We also found that glutamine deprivation in the culture media recapitulated the attenuated CTL proliferation along with the mTORC1 and c-Myc patterns generated under acidic conditions. However, while a 10-fold decrease in extracellular glutamine concentration resulted in the same intracellular levels measured at low pH, the impact on mTORC1 and c-Myc was only marginal. We observed that glutamine uptake by CTLs was lower under acidic conditions, at least in part due to competition with other amino acids for import. We also found that increased glutamine levels in the culture media was associated with lower intracellular levels of serine and threonine suggesting that low pH may disrupt substrate specificity and/or activity of ASCT2 (SLC1A5) which transports neutral amino acids inwardly only ^39, 40^. Taken together, our results suggest that lower levels of glutamine/glutamate/aspartate at low pH are consequences of either disturbed mTORC1/c-Myc status (especially considering c-Myc is a known regulator of glutamine metabolism) and/or of the activity of transporters involved in glutamine import/export.

Proline accumulation is a conserved stress response mechanism across many organisms including plants, bacteria, protozoa and marine invertebrates. In plants, for example, drought, cold, radiation, hyperosmolarity or pH changes drive proline build-up which can serves as an osmoprotectant and to quench reactive oxygen species (ROS) ^61, 62^. The role of elevated proline in CTLs, if any, is unclear. Indeed, the knockdown of P5CS (an enzyme involved in glutamate conversion to proline) prevented proline accumulation but it did not restore glutamine/glutamate/aspartate levels and there was no impact on mTORC1, c-Myc, CTL proliferation or viability.

We set out in this work with the aim of developing a combinatorial treatment and/or gene-engineering strategy allowing CTLs to resist the suppressive effects of extracellular acidity, a common feature of solid tumors. Indeed, while previous studies have demonstrated improved tumor control with proton pump inhibitors and bicarbonate ^19, 20^, such strategies may not be universally effective as tumor cells might also benefit from alkalization (*i.e.*, causing mTORC1 inhibition in tumor cells). Here, we have demonstrated that low pH causes profound disruptive changes in CTLs affecting responsiveness to IL-2, different intracellular signaling pathways, transcription factors, amino acid uptake and metabolism, which all together severely impair proliferation and effector functions. In the absence of directly restoring intracellular pH of CTLs, we conclude that it is unlikely that a simple solution exists for overcoming suppression caused by acidity in the TME.

## Acknowledgements

This work was generously supported by the Ludwig Institute for Cancer Research, the Swiss National Science Foundation (SNSF# 310030_204326 to MI), the ISREC Foundation, the Prostate Cancer Foundation, and the Carigest Foundation to GC and MI. We thank the Flow Cyometry Facility (Francisco Sala de Oyanguren), the Metabolomics Platform (Julijana Ivanisevic, Hector Gallart Ayala and Tony Teav) and the Cellular Imaging Facility (Florence Morgenthaler Grand) of the University of Lausanne for their expertise, Marjorie Decroux for technical help, and Patrick Reichenbach for useful scientific discussions. We also thank Prof. Chih Dang (The Johns Hopkins University School of Medicine) and Prof. Christoph Hess (University of Basel, University of Cambridge) for insightful scientific discussions.

## Contributions

R.V.S designed and directed the study, performed experiments, analyzed the data and wrote the paper. L.P. performed experiments. B.S. produced protein and plasmid constructs. I.C. carried out gene set enrichment analyses. G.C. directed the study. M.I. directed the study and wrote the paper.

## Materials and methods

### Mice

Mice were housed at the University of Lausanne (UNIL, Epalinges, Switzerland) animal facility. All *in vivo* experiments were conducted in accordance and approval from the Service of Consumer and Veterinary Affairs (SCAV) of the Canton of Vaud. Female C57BL/6 were purchased from Harlan (Harlan, Netherlands). OT-I (Charles River) and OT3 mice (kindly provided by Prof. Dietmar Zehn) carry a TCR transgene specific for ovalbumin. Homozygous OT-I x CRISPR/Cas9 mice were obtained upon crossing OT-I with B6J Rosa26-Cas9 mice (The Jackson Laboratory).

### Cell lines

The 293T and C1498 cell lines (ATCC) were grown in DMEM medium containing 4.5g/L glucose, sodium pyruvate, and glutamax (Gibco), supplemented with 10% heat-inactivated fetal bovine serum (SeraGold), 10mM HEPES, 50U/mL penicillin, and 50μg/mL streptomycin (Gibco). The CTLL-2 cell line (ATCC) was grown in the same medium as primary CTLs and was used to titrate the human IL-2 “switch” variant.

### Splenocyte isolation, CD8^+^ selection and CTL generation

Splenocytes were extracted from OT-I, OT3, or C57BL/6 mice. CD8^+^ T cells were purified by negative selection using the Miltenyi CD8^+^ T cell isolation kit (130-104-075), and were primed by culture on plates pre-coated with anti-CD3 antibodies (5μg/mL; clone 145-2C11, Biolegend) in the presence of anti-CD28 antibodies (1μg/mL; clone 37.51, Biolegend) and recombinant murine IL-2 (rmIL-2, 200IU/mL; Peprotech) at 37°C with 5% CO_2_. Every two or three days, cells were diluted, and expanded until nine to twelve days with fresh medium containing 200IU/mL rmIL-2.

PBMCs were extracted from buffy coats of healthy donors (Transfusion Interregionale CRS, Switzerland) using Ficoll-Paque Plus (Cytiva) density gradient centrifugation and CD8^+^ T cells were negatively selected using EasySep human CD8 T^+^ cell isolation kit (STEMCELL Technologies). CD8^+^ T cells were activated for three days using CD3/CD28-stimulation reagent (T Cell TransAct, Miltenyi Biotec), and expanded for four days using 2000IU/mL recombinant human IL-2 (Peprotech).

### CTL culture to analyze pH impact

Most of the experiments were carried out in DMEM, containing 1g/L glucose, sodium pyruvate and Glutamax (Gibco), supplemented with 10% heat-inactivated fetal bovine serum, non-essential amino acids (Gibco), penicillin-streptomycin (Gibco), and 50μM β-mercaptoethanol. When needed (e.g. for glutamine deprivation experiments), DMEM containing 1g/L glucose and sodium pyruvate (Gibco), supplemented with 10% dialyzed and heat-inactivated fetal bovine serum (Gibco), penicillin-streptomycin, phenol red (Sigma-Aldrich) and 50μM β-mercaptoethanol was used. Where specified, medium was complemented with 2mM glutamine (Bioconcept) or glutamax (Gibco). The medium was supplemented with HCl 1M and pre-incubated before use for 2 hours at 37°C, 5% CO_2_ in order to reach the desired pH (± 0.1 pH unit). In some experiments, the pH was restored to 7.4 with NaOH 1M. mTORC1 inhibitor (rapamycin), GSK-3β inhibitor (CHIR99021), Akt inhibitor (Akt1/2) and proteasome inhibitor (MG-132) were from Sigma-Aldrich. When used, DMSO was added as a vehicle in the control condition.

### Plasmid constructions

Mouse Rheb and mCherry-SEpHluorin (gift from Sergio Grinstein (Addgene plasmid # 32001; http://n2t.net/addgene:32001; RRID: Addgene_32001) ^33^ were codon optimized (Thermo Fisher Scientific) and cloned into a MSGV retroviral vector (gift from David Ott [Addgene plasmid # 64269; http://n2t.net/addgene:64269; RRID: Addgene_64269]) ^63^, straight after a Thy1.1 (CD90.1) reporter gene, a furin cleavage and a T2A ribosomal skipping sites. Using this plasmid, gene transcription upon virus integration is dependent on the LTR promoter constitutive activity. “miR Scramble” and “miR P5CS” are miR-30-based shRNA (Transomic Technologies) that were cloned in a modified version of the pQCXIP (Takara) plasmid under the control of the human U6 promoter along with the U6 snRNA leader sequence ^64^ (this plasmid is a self-inactivating retroviral vector modified to contain a Thy1.1 reporter gene under the control of the human phosphoglycerate kinase promoter, a woodchuck posttranscriptional regulatory element, and a bovine growth hormone polyA signal following the 3’ LTR). P5CS shRNA sequence was obtained with http://splashrna.mskcc.org/ online website prediction ^65^-the shRNA sequence used in this study (TCGACATGTAATTTCATTTCT) was the most potent out of three shRNAs assessed. The Scramble shRNA sequence (CAGGCAGAAGTATGCAAAGCA) is a negative control that does not match any sequence. sgRNA (obtained from http://chopchop.cbu.uib.no/ ^66^) against CD25 (AACCCCAACATCAGCAAGCG), Lkb1 (TCCTTAGCGCCCTACGTATA), Nprl2 (GCAGAGGCGGCCGTACCAAT), PRAS40 (ACGACATCGCACAGGCGCAC) or TSC2 (TCTCATACACTCGAGTGGCG) were cloned in a modified self-inactivated MSGV retroviral vector (Control sgRNA sequence: AAACCTAGCGTAGATTCGGC). Briefly, the vector was as follows: 5’LTR U3 promoter was replaced with a RSV promoter, the splicing donor site and gag sequence were removed, a U6 promoter, the sgRNA sequence including an optimized CRISPR tracrRNA sequence ^67^, poly-T terminator, a human PGK promoter, two copies of Thy1.1 split by furin and T2A sequences, and a WPRE sequence were added, the 3’LTR was modified in order to keep a minimal U3 sequence that is devoid of promoter activity and a bovine growth hormone polyA sequence was added after the U5. Constructs were validated by Sanger sequencing. The ecotropic packaging plasmid pCL-Eco was a gift from Inder Verma (Addgene plasmid # 12371; http://n2t.net/addgene:12371; RRID: Addgene_12371)^68^.

### Retrovirus production and transduction

293T were transfected using the calcium phosphate technique ^69^ with a 1:1 retroviral vector to packaging plasmid ratio. Two days post-transfection, supernatants were 0.45μm-filtered and concentrated with the retro-concentin virus precipitation solution (SBI). Retroviral content was titrated with the C1498 cell line in the presence of protamine sulfate (Sigma-Aldrich, P4020), based on CD90.1/Thy1.1 positivity by flow cytometry. Two days post-activation using the aforementioned protocol, CD8^+^ T cells were transduced (MOI of 2) with 10μg/cm^2^ of Retronectin (Takara) following manufacturer’s instructions. Resulting cells were expanded, as previously mentioned, with 200IU/mL rmIL-2. Three days post-infection, transduced cells were selected by magnetic separation (CELLection Biotin Binder Kit, ThermoFisher Scientific) via CD90.1/Thy1.1 surface labeling. Five to seven days post-transduction, purity (>90%) was confirmed by flow cytometry.

### Assessment of CTL proliferation and expansion

CTLs were labeled with the CellTrace Violet (CTV) Cell Proliferation Kit (5μM; Molecular Probes), re-activated with plates pre-coated with anti-CD3 antibodies (1μg/mL), and/or stimulated with various doses of IL-2, and cell division number was tracked by flow cytometry in the viable fraction. Cell expansion was obtained either by counting cell number by trypan blue exclusion using a Neubauer chamber, or by flow cytometry using Precision count beads (Biolegend).

### Cytotoxicity assay

As target cells, C1498 cells were labeled with 2.5 μM CTV for subsequent discrimination, and pulsed for one hour with different concentrations of SIINFEKL peptide (Protein and Peptide Chemistry Facility of the University of Lausanne). Resulting cells were co-cultured with CTLs for four hours at various CTL-to-tumor ratios. C1498 cell death was quantified by flow cytometry with the live/dead fixable near-IR dead cell stain kit (Molecular Probes). CTL-mediated lysis (%) was calculated as follows: (%CTL-induced cell death - %spontaneous cell death)/(100 - %spontaneous cell death). Antigen-specific lysis (%) was calculated as: (%CTL-induced cell death of Ag-pulsed C1498 - %CTL-induced cell death of unpulsed C1498)/(100 - %CTL-induced cell death of unpulsed C1498).

### Cytokine secretion analyses

Cytokine content (IFN-γ, IL-2 and TNF) from supernatant of re-activated CTLs was determined using the mouse ELISA MAX Sets from Biolegend according to the manufacturer’s instructions.

### Intracellular pH determination by confocal microscopy

SEpHluorin-transduced CTLs were pre-stained with Hoechst 34580 (Thermo Fisher Scientific) to stain nuclei. For standard curves, CTLs were seeded on poly-L-Lysine-coated Labtek I culture chamber slides (Nunc, 055083) in high K^+^ buffer (145mM KCl, 1mM MgCl_2_, 0.5mM EGTA, 10mM MES, 10mM HEPES) containing nigericin and valinomycin (10μM each, Thermo Fisher Scientific), and adjusted to pH7.8, pH7.3, pH6.8 and pH6.3 with NaOH. This protocol allows to equilibrate intracellular pH to extracellular pH. In parallel, CTLs were seeded on poly-L-Lysine-coated Labtek I culture chamber slides at pH7.4 or pH6.6 in bicarbonate and phenol-free media containing 15mM HEPES and 15mM MES in the presence of rmIL-2 (200 IU/mL). CTLs were imaged by confocal microscopy each 40 minutes.

Confocal fluorescence images were acquired with a Zeiss LSM800 confocal microscope using a 40x objective without immersion. Experiments were carried out at 37°C in a humidified atmosphere without CO_2_ using an incubation chamber. At least 4 positions per well were acquired. Resulting images were processed with *in-house* ImageJ/Fiji macros in order to obtain nucleus-free or cytoplasm-free pictures and fluorescence values. At least 100 cells were analyzed per condition.

### Cell cycle analysis

Briefly, CTLs were fixed and permeabilized with 70% Ethanol, depleted from endogenous RNA using RNAse A (Sigma-Aldrich), and DNA was stained using propidium iodide (ThermoFisher Scientific). Resulting cells were analyzed by flow cytometry.

### IL-2R complexes quantification and binding capacities of IL-2 to IL-2R

rmIL-2 was biotinylated using the EZ-Link Sulfo-NHS-LC-Biotin kit (ThermoFisher Scientific) at a 100 biotin to 1 IL-2 molar ratio, as previously described by De Jong et al ^70^, following manufacturer’s instructions, and was serially filtered with an AMICON centrifugal unit (Merck) to remove unbound biotin. Following culture of CTLs under various pH, IL-2R complexes were determined by staining the cells with the biotinylated-rmIL-2 (equivalent to 200IU/mL) at 4°C for 30 minutes. Cells were stained with PE-conjugated streptavidin (Biolegend), the signal was further amplified with another round of staining with biotinylated anti-streptavidin (Vector Laboratories) and PE-conjugated streptavidin, and analyzed by flow cytometry. The impact of pH on binding capacities of IL-2 to IL-2R was determined by incubating biotinylated-rmIL-2 (200IU/mL) with CTLs under various pH in PBS at 4°C for 30 minutes. IL-2 binding was determined following the same aforementioned protocol.

### Production and purification of human IL-2 “switch”

Rosetta pRAREII (Novagen) were transformed with the pET-22b plasmid encoding the IL-2 “switch” sequence ^29^ and protein was extracted from inclusion bodies. Non-aggregated protein was enriched by preparative size-exclusion column chromatography (Superdex75) followed by Amicon filtration.

### Staining for flow cytometry

For all the experiments, but intracellular pH detection, the buffer used was composed of PBS, 1% BSA, 0.1% NaN_3_, and the staining procedure included the use of a live/dead fixable cell stain kit (Molecular Probes) to analyze viable cells. For surface marker expression, cells were stained with CD8α-AF488 (100723), -PECy7 (100722), or –BV421 (100738), CD25-PECy7 or APC (102016, 102012), CD44-PECy7 (103030), CD62L-BV570 (104433), CD90.1/Thy1.1-APC or BV421 (202526, 202529), or – biotin (202510), CD132-APC (132308), or GrB-AF647 (515406) antibodies from Biolegend, or CD122-BV421 (564925) antibody from BD Biosciences, or appropriate isotype-matched controls. For apoptosis determination, cells were stained with AnnexinV-FITC (BD Biosciences) and the live/dead fixable near-IR dead cell stain kit. Apoptosis was analyzed amongst viable (live/dead marker^-^) cells, and AnnexinV^+^ cells were considered as apoptotic. Samples were run using a Gallios, a CytoFlex (Beckman Coulter) or a BD FACSCanto flow cytometer. Analyses were carried out on singlets using FlowJo software, and marker expression was calculated from the Median Fluorescence Intensity (MFI) by calculating the ratio MFI (rMFI) as: MFI marker/MFI isotype (or, MFI unstained in case isotype was not available, *e.g.* for biotinylated-IL-2 staining, MFI biot-IL-2/MFI w/o biot-IL-2). Then, the background was removed (rMFI-1), and the result was normalized to the “control” condition (*i.e.* the “pH7.4” condition, which is considered to be equivalent to 100%), such as: normalized rMFI (%) = (rMFI[test]-1) x100/(rMFI[control]-1).

### mRNA expression analyses by real-time quantitative PCR

DNA-free RNA was extracted from dry pellets using the Qiagen RNeasy kit and DNAse Set according to the manufacturer’s instructions. Equal amounts of RNAs were used to synthesize cDNA with the oligodT primer from the Primescript first strand cDNA synthesis kit (Takara Bio). *Myc* (forward: TTGATGTGGTGTCTGTGGAGAAGAG; reverse: CGTAGTTGTGCTGGTGAGTGGA) expression was analyzed by normalization with *Actb* (forward: CTAAGGCCAACCGTGAAAAGAT; reverse: CACAGCCTGGATGGCTACGT) through real-time quantitative PCR (SDS 7900 HT instrument; Applied Biosystems) using the Kapa Sybr Fast mix (Roche). The following parameters were used: 95°C for 3 min, and 40 cycles of 95°C 3 sec, 60°C 30 sec and 72°C 1 sec. Each reaction was performed in duplicate. The 2^ΔCt^ method was used to normalize and linearize the results, but was adjusted to the observed primer efficacy (*i.e. Myc*: 1.80; *Actb*: 1.98).

### RNA sequencing

RNA quality was assessed on a Fragment Analyzer (Agilent Technologies) and all RNAs had a RQN between 7.5 and 10. RNA-seq libraries were prepared using 350 ng of total RNA and the Illumina TruSeq Stranded mRNA reagents (Illumina) on a Sciclone liquid handling robot (PerkinElmer) using a PerkinElmer-developed automated script. Cluster generation was performed with the resulting libraries using the Illumina HiSeq 2500 SR Cluster Kit v4 reagents and sequenced on the Illumina HiSeq 2500 using TruSeq SBS Kit v4 reagents. Sequencing data were demultiplexed using the bcl2fastq Conversion Software (v. 2.20, Illumina).

### Immunoblotting

Unless specified, prior to the assay cells were starved for one hour and a half by culture at 37°C in cytokine- and serum-free medium. For the assays, the resulting cells were cultured (1-2 million cells per mL) with rmIL-2 at various doses for different times. Cells were lysed in RIPA buffer supplemented with Halt phosphate/protease inhibitors (ThermoFisher Scientific) and were boiled at 97°C for 10 minutes with Bolt LDS sample buffer and reducing agent (ThermoFisher Scientific). Protein samples were separated by SDS-PAGE and transferred to PVDF membranes using the iBlot2 system (Thermo Fisher Scientific). Antibody staining of the different molecules of interest was carried out according to the manufacturer’s instructions. Antibodies against p-Akt (4060 and 13038), Akt (4691), p-AMPKα (2535), AMPKα (5831), p-AMPKβ (4186), AMPKβ (4150), p-CREB (9198), p-c-MycS62 (13748), c-Myc (5605), p-eIF2α (3398), p-ERK (4377), ERK (4695), p-FoxO1/3 (9464), p-GSK-3β (9323), p-JAK1 (74129), JAK1 (3344), p-JAK3 (5031), JAK3 (8863), LKB1 (3047), NPRL2 (37344) p-PLCγ1 (2821), PLCγ1 (2822), PRAS40 (2691), p-PRAS40 (2997), p-p38 (4511), p62 (5114), p-p70S6K (9205), p70S6K (9202), Rheb (13879), p-STAT5 (9359), p-S6 (4858 and 5364), p-SLP-76 (92711), SLP-76 (4958), Tuberin/TSC2 (4308), p-ULK1 (6888), ULK1 (8054), 4E-BP1 (9644) and p-4E-BP1 (2855 and 9451) were from Cell Signaling Technology. Antibodies against CREB (A301-669A) and STAT5 (A303-494A) were from Bethyl Laboratories. Anti-p-c-MycT58 (ab28842) antibody was from Abcam, anti-FoxO (MA5-32114), HRP-conjugated anti-mouse IgG (A16072) and HRP-conjugated anti-rabbit IgG (31458) antibodies were from ThermoFisher Scientific, while those against β-Actin (sc-47778), CREM (sc-390426), GAPDH (sc-32233) and S6 (sc-74459) were from Santa Cruz. Antibody against ALDH18A1/P5CS (HP008333) was from Sigma-Aldrich. Images were acquired with a western blot imager (Fusion, Vilber Lourmat) and protein levels were quantified using the ImageJ software by analyzing pixel intensity. Membranes were stripped (Tris 62.5mM pH6.7, SDS 2%, 0.1M β-mercaptoethanol, 30 minutes at 50°C) in order to stain the same membrane for different proteins. Quantification was carried out using the ImageJ/Fiji software. For most of the molecules, phosphorylation status was calculated by dividing the signal of the phosphorylated protein by the signal of the matched total protein (except phosphorylation of eIF2α, GSK-3β, and p38, for which the signal was divided by the β-actin signal), which were acquired from the same membrane. Total levels of the other molecules assessed were calculated by dividing their signal to the β-actin signal. Results were normalized to the control condition (*i.e.* the “pH7.4 + IL-2” condition yields 100%).

### Glutamine isotopic trace analysis

CTLs were pre-cultured 2 hours in the absence of serum, IL-2 and glutamine a 37°C. Upon culture for 4 hours at 37°C in CTL medium containing dialyzed serum, IL-2 and ^13^C-glutamine (2mM – Sigma Aldrich, 605166), cell pellets were stored at -80°C. CTL lysates (2 million cells) were extracted and homogenized by the addition of 500 µL of MeOH 80% and ceramic beads, in the air-cooled Cryolys Precellys Homogenizer (2 x 20 seconds at 10000 rpm, Bertin Technologies, Rockville, MD, US). Homogenized extracts were centrifuged for 15 minutes at 21000 g and 4°C, and the resulting supernatant was collected and evaporated to dryness in a vacuum concentrator (LabConco, Missouri, US). Dried sample extracts were re-suspended in MeOH:H_2_O (4:1, v/v) prior to LC-HRMS analysis according to the total protein content (to normalize for sample amount).

Cell extracts were analyzed by Hydrophilic Interaction Liquid Chromatography coupled to high resolution mass spectrometry (HILIC - HRMS) in negative ionization mode using a 6550 Quadrupole Time-of-Flight (Q-TOF) system interfaced with 1290 UHPLC system (Agilent Technologies) as previously described ^71^.

Raw LC-MS files were processed in Profinder B.08.00 software (Agilent Technologies) using the targeted data mining in isotopologue extraction mode. The metabolite identification was based on accurate mass and retention time matching against an *in-house* database. The Extracted Ion Chromatogram areas (EICs) of each isotopologue (M+0, M+1, M+2, M+3,…) were corrected for natural isotope abundance ^72^ and the label incorporation or ^13^C enrichment was calculated based on relative isotopologue abundance (in %), in each one of two analyzed conditions ^73^.

In addition, pooled QC samples (representative of the entire sample set) were analyzed periodically (every 4 samples) throughout the overall analytical run in order to assess the quality of the data, correct the signal intensity drift and remove the peaks with poor reproducibility. In addition, a series of diluted quality controls (dQC) were prepared by dilution with methanol: 100% QC, 50%QC, 25%QC, 12.5%QC and 6.25%QC and analyzed at the beginning and at the end of the sample batch. This QC dilution series served as a linearity filter to remove the features that do not respond linearly or correlation with dilution factor is < 0.65.

### Amino acid quantification

CTL lysates (1-2 million cells) were extracted and homogenized by the addition of 750 µL of MeOH 80% and ceramic beads, in the air-cooled Cryolys Precellys Homogenizer (2 x 20 seconds at 10000 rpm, Bertin Technologies, Rockville, MD, US). Homogenized extracts were centrifuged for 15 minutes at 21000 g and 4°C. The 50 µL of extract were mixed with 250 µL of Methanol containing isotopic labeled internal standards. Sample extracts were vortexed and centrifuged (15 min, 2700g at 4°C) and the resulting supernatant was collected and injected into the HILIC-HRMS (Q Exactive Focus instrument interfaced with the a HESI source) (Thermo Fisher Scientific) for amino acid quantification ^74^.

### Total protein quantification

After metabolite extraction with organic solvents (MeOH) the obtained protein pellets were evaporated and lysed in 20 mM Tris-HCl (pH 7.5), 4M guanidine hydrochloride, 150 mM NaCl, 1 mM Na_2_EDTA, 1 mM EGTA, 1% Triton, 2.5 mM sodium pyrophosphate, 1 mM β-glycerophosphate, 1 mM Na_3_VO4, 1 µg/ml leupeptin using the Cryolys Precellys 24 sample Homogenizer (2 x 20 seconds at 10000 rpm, Bertin Technologies, Rockville, MD, US) with ceramic beads. BCA Protein Assay Kit (Thermo Scientific, Masschusetts, US) was used to measure (A562nm) total protein concentration (Hidex, Turku, Finland).

### Amino acid uptake assay

CTLs were incubated at 5 million per mL in HBSS buffer (Invitrogen) at various pH for 20 minutes at room temperature in the presence of 10μCi/mL of tritiated glutamine (equivalent to 165nM; Hartmann Analytic. Alternatively, CTLs were pre-incubated (37°C, 20 minutes) under various pH in CTL medium containing IL-2 with dialyzed serum without glutamine and were further incubated for 20 minutes in the presence of 10μCi/mL of tritiated glutamine. Cells were washed, lysed with 0.3% final SDS, and radioactivity incorporation was determined with a scintillation counter (PerkinElmer) after MicroScint 40 (PerkinElmer) addition.

### In vitro mTORC1 kinase activity

Kinase activity of recombinant mTORC1 complexes (human mTOR/RAPTOR/MLST8, Sigma-Aldrich) was assessed by determining phosphorylation status of recombinant human 4E-BP1 (Abcam). Kinase assay was carried out with 20mM HEPES, 20mM MES, 10mM MgCl_2_ and 50mM KCl. 10x reaction buffer were prepared by adding NaOH to reach pH7.4 and pH6.6. mTORC1, 4E-BP1 and reaction buffer were mixed and preincubated at room temperature for 20 minutes. Upon addition of 200μM ATP, samples were incubated at 30°C for 10 minutes. Thereafter, samples were boiled at 97°C for 10 minutes with Bolt LDS sample buffer and reducing agent, and were separated by SDS-PAGE and transferred to PVDF membranes. Staining was carried out according to the beforehand described section for Western blot.

### Statistical analyses

Statistical significance was evaluated using the two-tailed paired student’s t-test for comparison of two groups, or a one- or two-way ANOVA with a Turkey post-hoc test for comparison of multiple groups using the GraphPad Software (Prism). For correlation curves, linear and non-linear correlation curves were modelled with GraphPad according to plot patterns.

---

**Extended Data Fig. 1:**
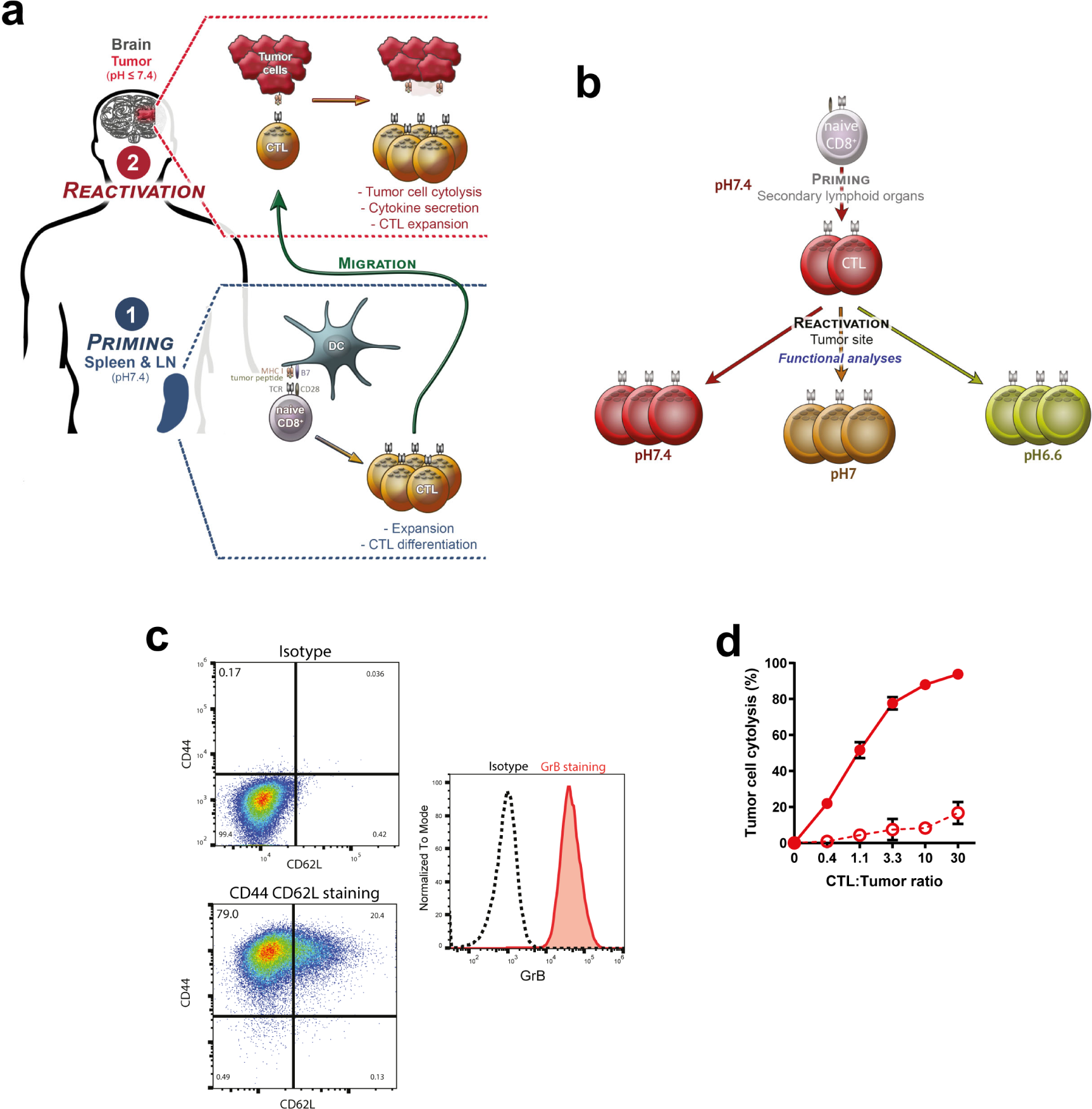
Rationale, methodology, and validation of the *in vitro*-designed model to study the impact of low pH on the anti-tumor CD8^+^ T cell response. **a,** Illustration of the main steps and functions of anti-tumor CD8^+^ T cell migrating to the tumor microenvironment. Previously differentiated anti-tumor CD8^+^ T cells (i.e. effector cytotoxic T lymphocytes; CTLs) encounter acidic conditions at the tumor site when they exert their activities. **b,** Scheme depicting the in *vitro* methodology used to de-convolute the impact of pH on CTLs function in tumors. **c,** Effector/memory phenotype of primed and expanded OT-I CD8^+^ T cells used in this study. Naïve OT-I CD8^+^ T cells were activated for two days with pre-coated anti-CD3 and soluble anti-CD28 antibodies, and expanded for further ten days in the presence of murine IL-2. Dot plots show one representative experiment of background staining (“Isotype”, top graph), and of CD44 CD62L staining (bottom graph) obtained by flow cytometry. Histograms show one representative experiment of background staining (“Isotype”, black dashed line), and of Granzyme B staining (red solid line) obtained by flow cytometry. Almost 80% of cells are effectors (CD44^+^ CD62L^-^), while all express Granzyme B to relatively high levels. **d,** Effector generation protocol gives efficient killing capacities of the resulting OT-I. Varying numbers of OT-I CTLs were co-cultured for 4 hours and a half with C1498 tumor cells pulsed (solid lines and symbols), or not (dashed lines and empty symbols), with 1μM of antigen (minimal ovalbumin peptide epitope, SIINFEKL). Results show the mean percentage of tumor cell lysis ± SD of at least three (or two for 0.4 and 1.1 CTL:Tumor ratios) biological replicates from at least two independent experiments.

**Extended Data Fig. 2:**
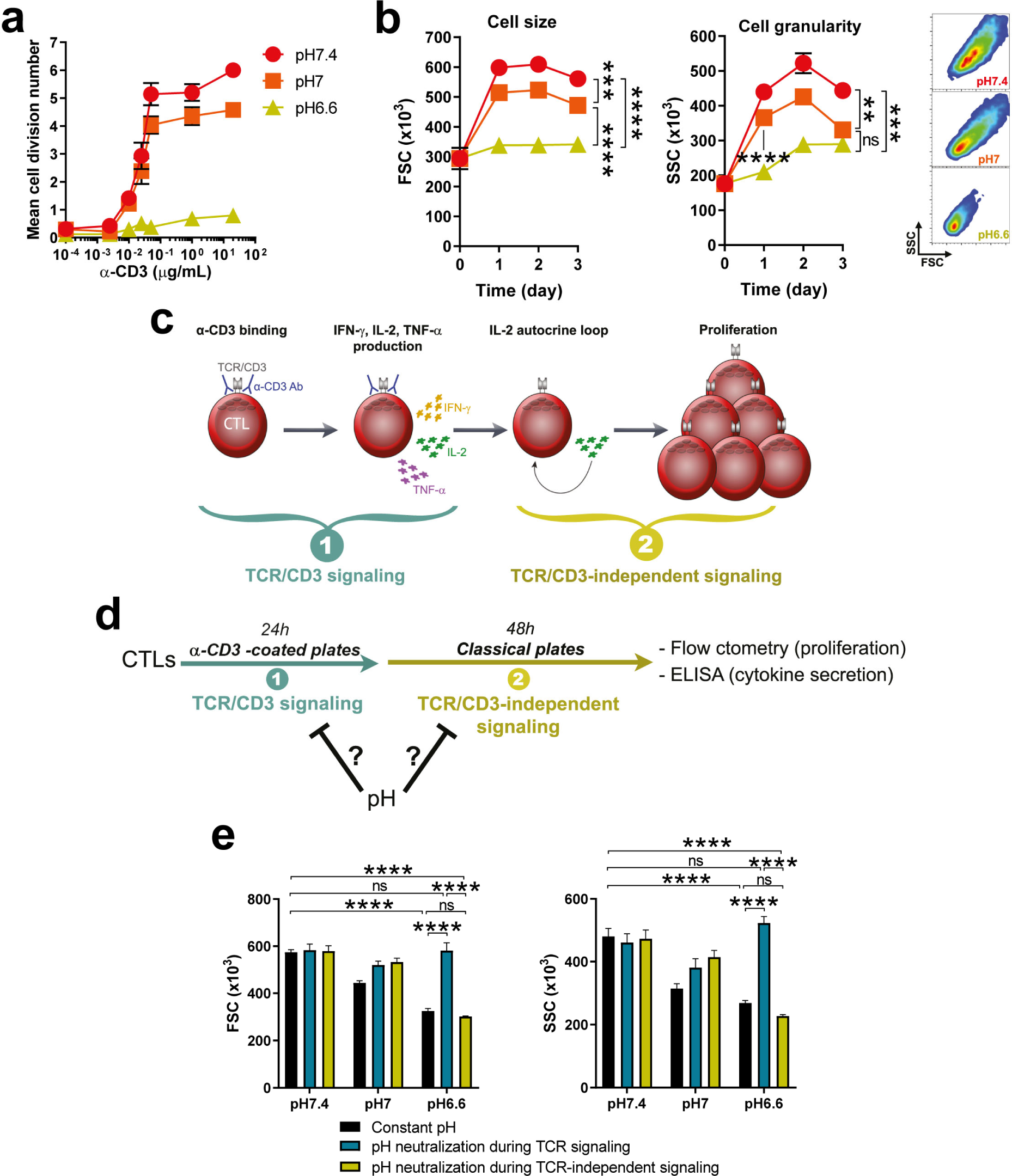
Impact of acidity on CTL size, granularity and proliferation in response to anti-CD3 stimulation. **a,** Impact of pH on CTL proliferation following re-activation with increasing doses of anti-CD3. OT-I CTLs were cultured for three days in the presence of various doses of an agonistic anti-CD3 antibody (pre-coated plates) at different pH (pH7.4, red circles, pH7, orange squares, or pH6.6, lime triangles). Line graph show the mean cell division number ± SEM of at least three biological replicates from at least two independent experiments. **b,** Low pH prevents the increase of CTL size and granularity following anti-CD3 re-activation. OT-I CTLs were cultured for one, two, and three days in the presence of an agonistic anti-CD3 antibody (1μg/mL pre-coated plates) at different pH (pH7.4, red circles, pH7, orange squares, or pH6.6, lime triangles). Bar graph shows the mean cell size or granularity ± SEM of at least four biological replicates from at least two independent experiments. Density plots shows one representative experiment obtained by flow cytometry one day post re-stimulation. ns: not statistically significant, **p<0.01, ***p<0.001, ****p<0.0001 (Student’s paired *t*-test). **c,** Cartoon depicting the two main steps involved in cytokine secretion and proliferation following anti-CD3 re-activation of CTLs. The TCR/CD3–independent signaling allowing CTL proliferation is mostly mediated by IL-2/IL-2R signaling. **d,** Design of the experiments in order to analyze at which step pH impacts cytokine secretion and proliferation during anti-CD3 –mediated CTL re-activation. pH was neutralized or acidified at step 1, or 2, in order to know whether cytokine secretion or proliferation can be restored. **e,** Acidity influences cell size and granularity of CTLs during TCR signaling. OT-I CTLs were cultured for one day in anti-CD3 –coated plates (TCR signaling step). The resulting cells and supernatants were transferred to an anti-CD3 –free plate and cultured for two further days (TCR-independent signaling step). Cells were either: cultured during the TCR signaling step and the TCR-independent signaling at constant pH (either pH7.4, pH7, pH6.6 – black bars, “Constant pH”), cultured at pH7.4 during the TCR signaling step then at various pH during the TCR-independent signaling step (“pH neutralization during TCR signaling” – teal bars), or cultured at various pH during TCR signaling step then at pH7.4 during the TCR-independent signaling step (“pH neutralization during TCR-independent signaling” – lime bars). Bar graph shows the mean (**c**) cell size or (**d**) granularity +SEM of four biological replicates from two independent experiments. ns: not statistically significant, ****p<0.0001 (one-way repeated measures ANOVA, Turkey post-hoc test).

**Extended Data Fig. 3:**
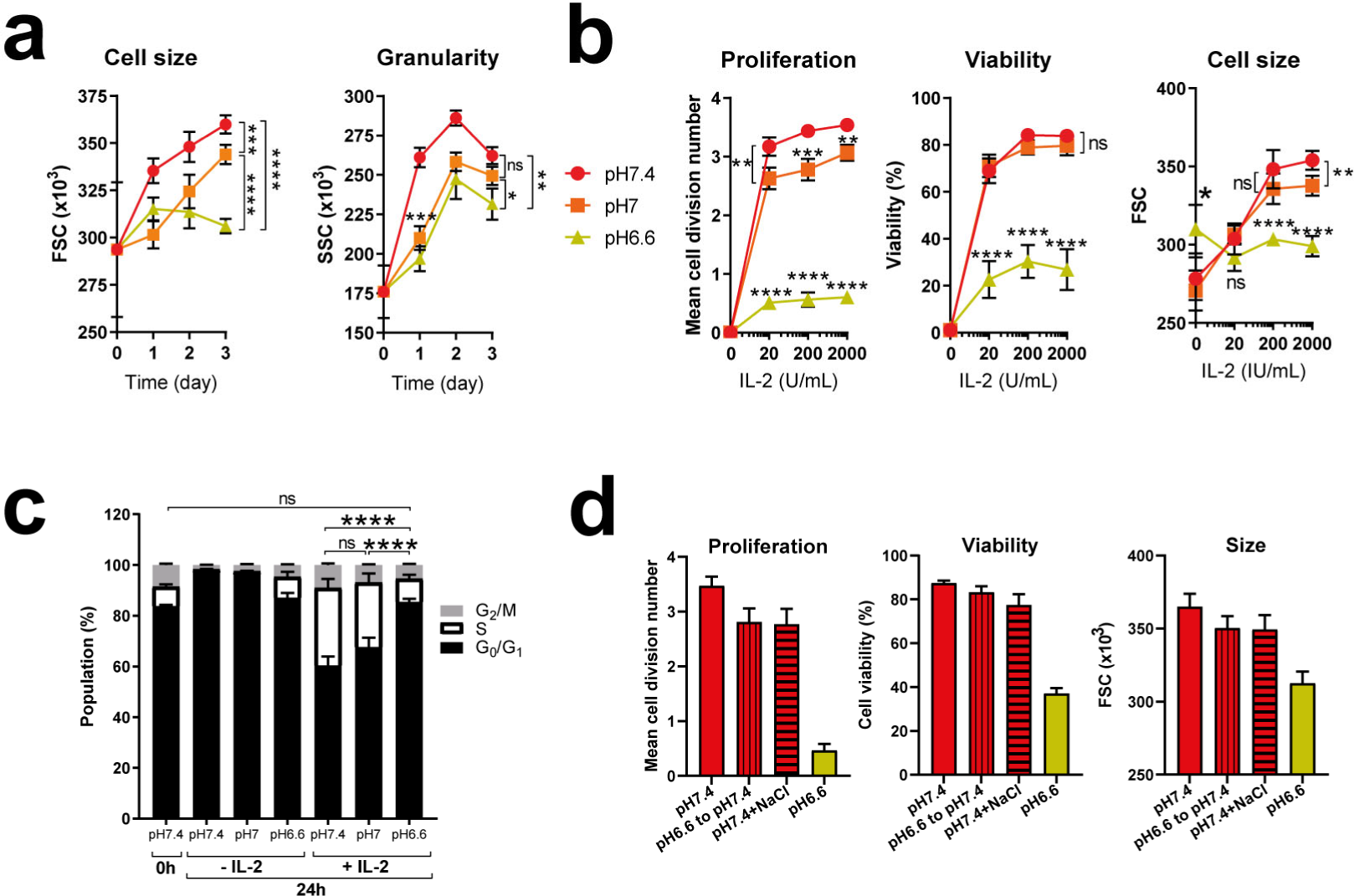
Acidity lowers IL-2-dependent response of CTLs. **a,** Time-course of the pH impact on the cell size and granularity of CTLs to IL-2. OT-I CTLs were cultured for one, two, or three days at three different pH (pH7.4, red circles, pH7, orange squares, or pH6.6, lime triangles) in the presence of exogenous murine IL-2 (200 IU/mL). Results show the mean cell size, or granularity, ± SEM of at least three biological replicates from at least two independent experiments. ns: not significant, *p<0.05, **p<0.01, ***p<0.001, ****p<0.0001 (Student’s paired *t*-test). **b,** Impact of pH on the dose-response of CTLs to IL-2. OT-I CTLs were cultured for three days at three different pH (pH7.4, red circles, pH7, orange squares, or pH6.6, lime triangles) in the presence of various doses of murine IL-2. Results show the mean cell division number, viability or cell size ± SEM of at least seven biological replicates from at least three independent experiments. ns: not significant, *p<0.05, **p<0.01, ***p<0.001, ****p<0.0001 (Student’s paired *t*-test). **c,** Impact of pH on cell cycle phases of CTLs in response to IL-2. OT-I CTLs were cultured for one day at three different pH in the presence, or absence, of exogenous murine IL-2 (200 IU/mL). Results show the mean proportion of cells in G_0_/G_1_ (black solid bars), S (black empty bars), or G_2_/M (grey solid bars) phases as evidenced by propidium iodide staining +SEM of four biological replicates from two independent experiments. ns: not significant, ****p<0.0001 (one-way repeated measures ANOVA, Turkey post-hoc test): calculated for the S phase. **d,** Impact of acidity on CTL response to IL-2 is not due to precipitation/inactivation of molecules in the medium or to increased osmolarity. OT-I CTLs were cultured for three days at pH7.4 or pH6.6 in the presence of exogenous murine IL-2 (200 IU/mL). The conditions “pH6.6 to pH7.4” refers to the use of medium that was beforehand acidified at pH6.6, incubated for two hours at 37°C and neutralized with NaOH to pH7.4, while “NaCl” means that CTLs were cultured at pH7.4 in the presence of the same amounts of NaCl than the HCl concentrations (around 31 mM) used to reach pH6.6. Results show the mean cell division number, viability or cell size +SEM of five biological replicates from two independent experiments.

**Extended Data Fig. 4:**
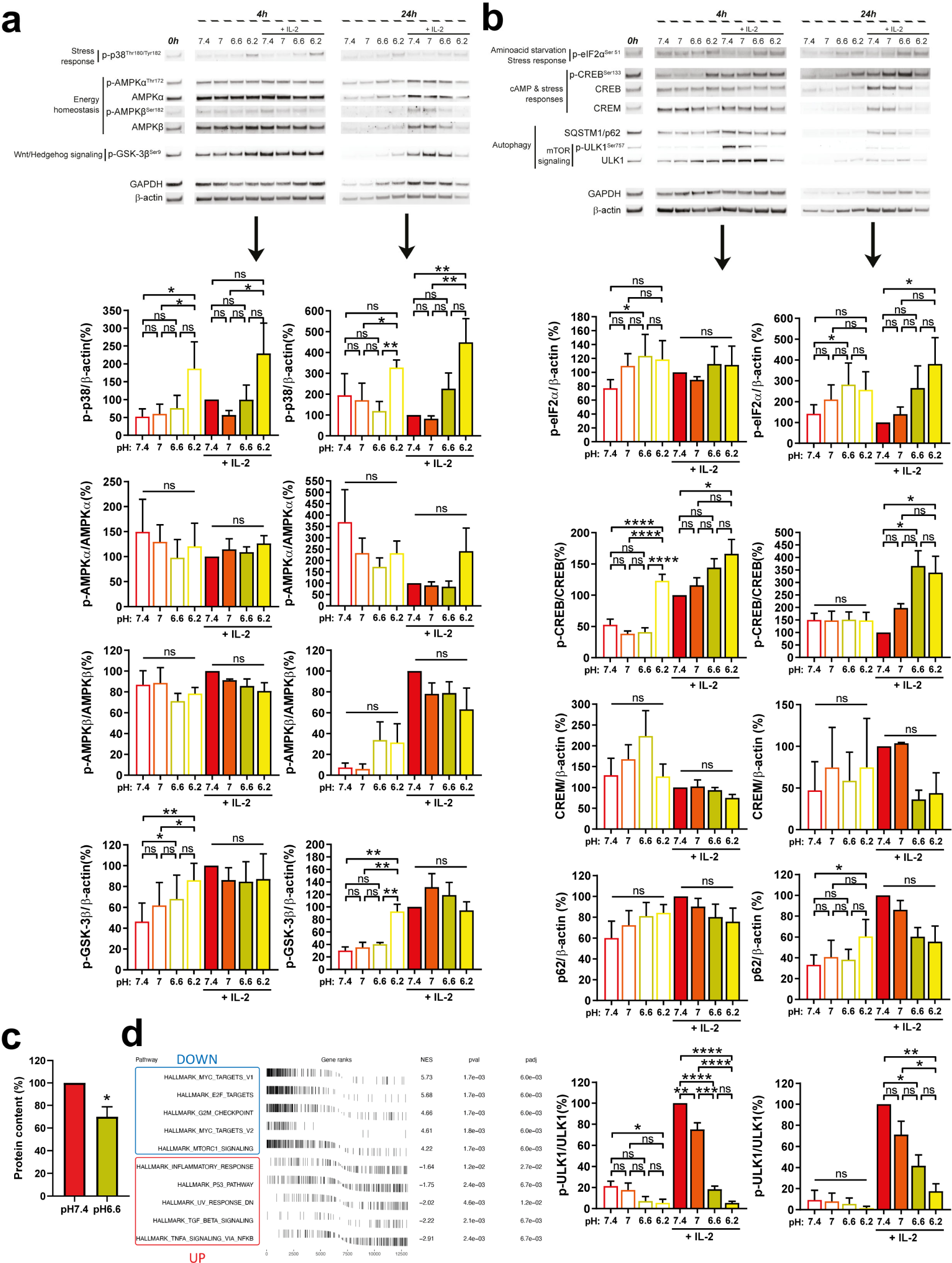
Impact of pH on pathways that may act on mTORC1 and c-Myc. **a,b,** Impact of pH on pathways that could lead to mTORC1 and c-Myc modulations. OT-I CTLs were cultured at various pH in the presence, or the absence of exogenous murine IL-2 (200 IU/mL) for 4 or 24 hours. One representative western blot is shown. Bar graphs show the mean phosphorylation status, or total levels, of the indicated molecule relative to the total protein of interest, or to β-actin, normalized to the condition “pH7.4 + IL-2” ± SEM of at least three biological replicates from at least two independent experiments. ns: not significant, *p<0.05, **p<0.01, ***p<0.001, ****p<0.0001 (one-way repeated measures ANOVA, Turkey post-hoc test). **c,** Acidity limits protein content. OT-I CTLs were cultured for four hours in the presence of exogenous IL-2 (200 IU/mL), at pH7.4 or pH6.6. Bar graph shows mean protein content, normalized to the condition “pH7.4”, +SEM of four biological replicates from two independent experiments. *p<0.05 (Student’s *t*-test). **d,** Acidity blunts c-Myc targets expression. OT-I CTLs were cultured for twenty-four hours in the presence of exogenous IL-2 (200 IU/mL), at pH7.4 or pH6.6. Results displays GSEA from RNAseq data from five biological experiments from two independent experiments.

**Extended Data Fig. 5:**
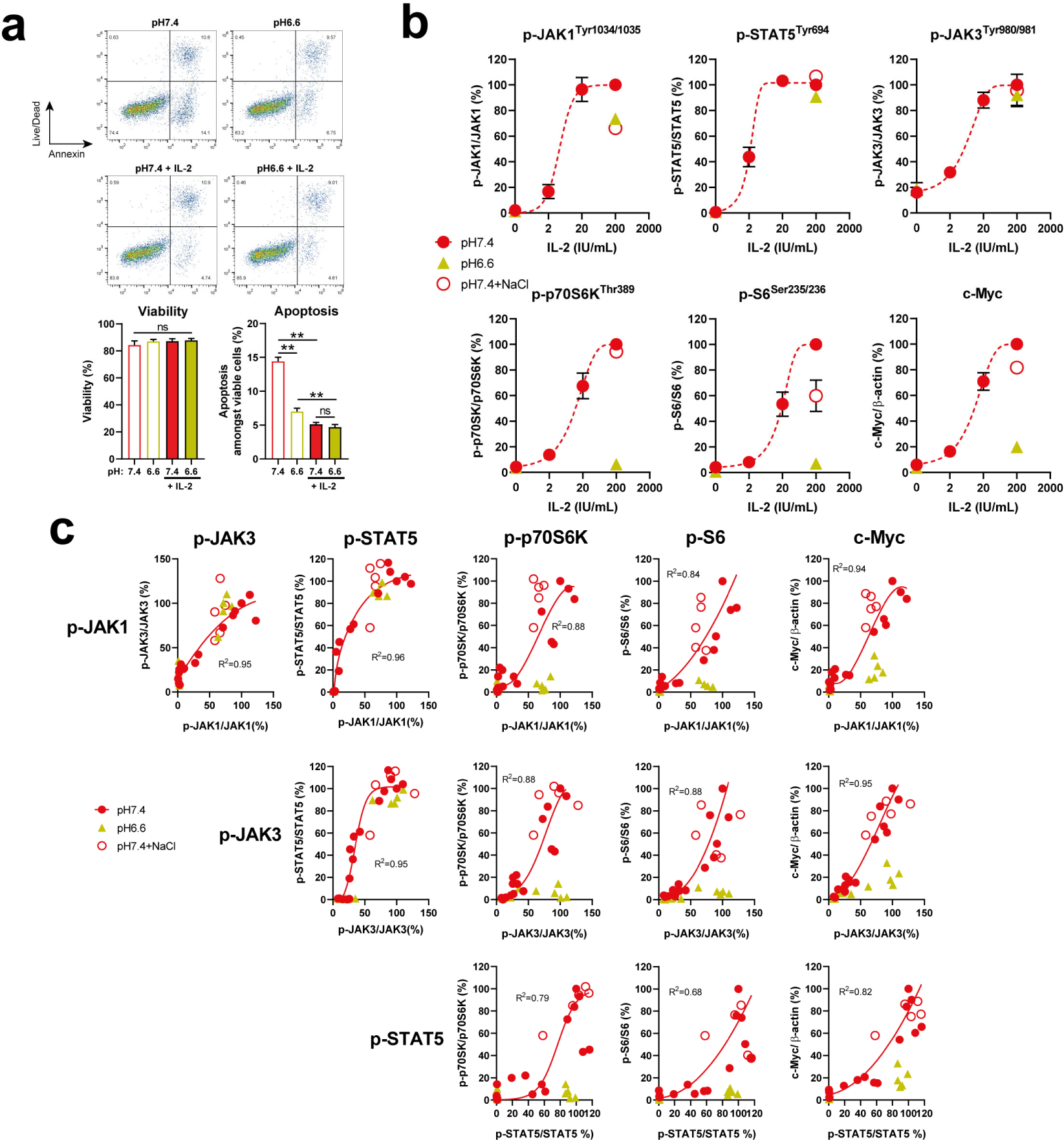
The impact of acidity on mTORC1 and c-Myc is not linked to a disturbance in the activation of first IL-2R signaling transducers in polyclonal CTLs. **a,** Acidity does not increase apoptosis or cell death of CTLs upon a four-hour treatment. OT-I CTLs were cultured at various pH in the presence, or the absence of exogenous murine IL-2 (200 IU/mL) for 4 hours. Representative dot plots show Annexin and Live/Dead staining obtained by flow cytometry. Bar graph shows percent cell viability (Live/Dead negative fraction) or apoptosis amongst viable cells (Annexin positive amongst Live/Dead negative fraction) + SEM of four biological replicates from two independent experiments. **b,** Dose-dependent activation of the first signaling transducers, mTORC1 pathway and accumulation of c-Myc. C57BL/6 CTLs were cultured at pH7.4 (red circles and dashed lines) in the presence of various concentrations of exogenous murine IL-2 for 4 hours. In parallel, CTLs were cultured at pH7.4 with 31mM NaCl (red empty circles) or at pH6.6 (lime triangles) with exogenous murine IL-2 (200 IU/mL). Results show the mean phosphorylation status, or total levels, of the indicated molecule relative to the total protein of interest, or to β-actin, normalized to the condition “pH7.4 + 200 IU/mL IL-2” ± SEM of at least four biological replicates from two independent experiments. **c,** Correlation between the activation of first signaling transducers and mTORC1 pathway targets activation, or c-Myc levels. C57BL/6 CTLs were cultured at pH7.4 (red circles) in the presence of various concentrations of exogenous murine IL-2 for 4 hours. In parallel, CTLs were cultured at pH7.4 with 31mM NaCl (red empty circles) or at pH6.6 (lime triangles) with exogenous murine IL-2 (200 IU/mL). Correlation curves, and the associated R^2^, were calculated for the condition “pH7.4” (red solid lines). Results show individual values obtained from **a,**.

**Extended Data Fig. 6:**
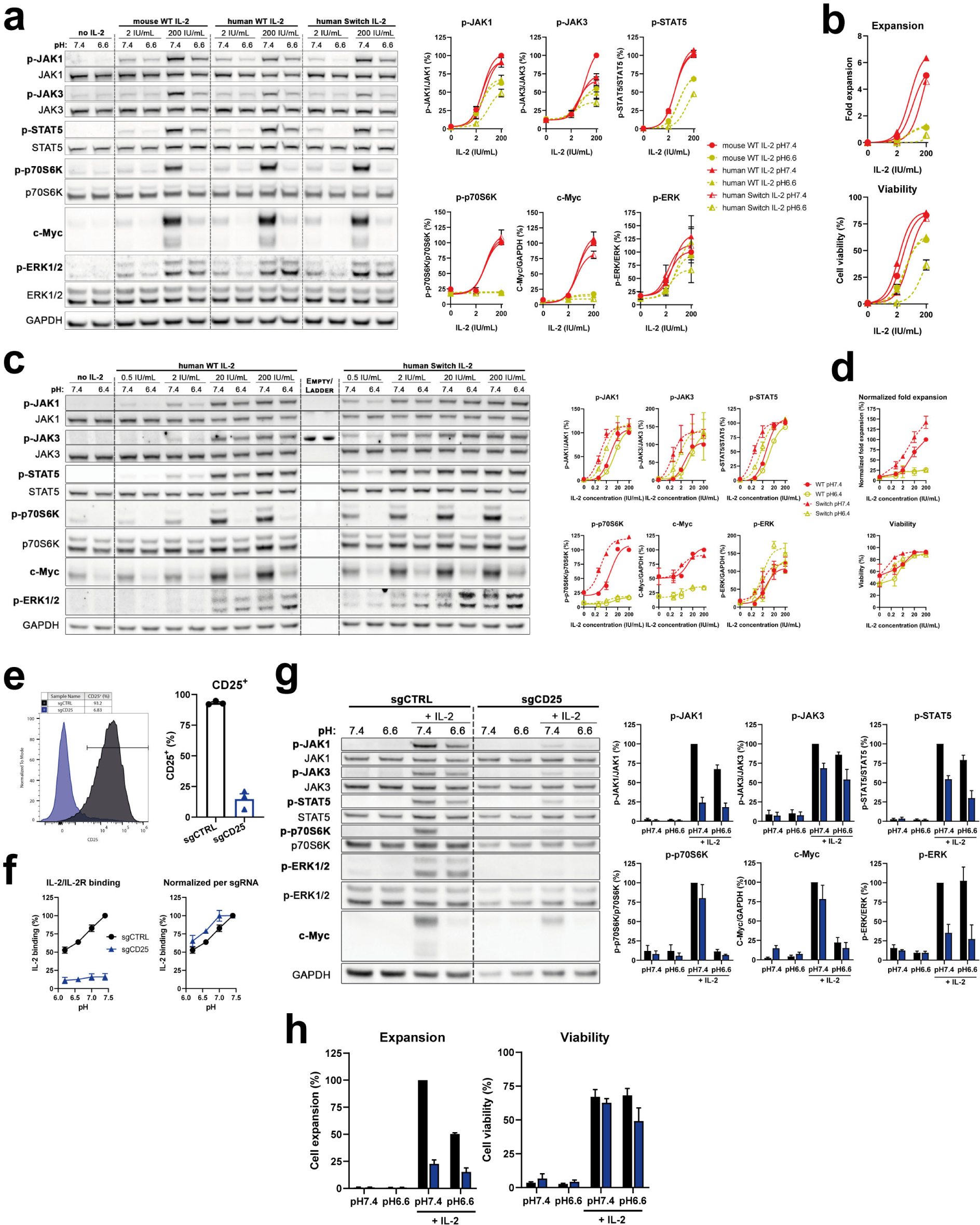
The Switch IL-2 mutant does not improve mTORC1, c-Myc and cell expansion at low pH. **a,** Impact of the IL-2 “Switch” cytokine on IL-2R signaling. OT-I CTLs were cultured for four hours in the presence, or absence, of wild type mouse IL-2 (circles), wild type human IL-2 (solid triangles) or human “Switch” IL-2 (empty traingles) at pH7.4 (red and solid lines) or pH6.6 (lime and dashed lines). Blots of one representative experiment are shown. Line graphs shows the mean ± SEM of three biological replicates from two independent experiments. **b,** Impact of the IL-2 “Switch” cytokine on expansion and viability. OT-I CTLs were cultured for three days in the presence, or absence, of wild type mouse IL-2 (circles), wild type human IL-2 (solid triangles) or human “Switch” IL-2 (empty traingles) at pH7.4 (red and solid lines) or pH6.6 (lime and dashed lines). Line graphs shows the mean ± SEM of three biological replicates from two independent experiments. **c,** Impact of the IL-2 “Switch” cytokine on IL-2R signaling in human CD8^+^ T cells. Human CD8^+^ T cells were cultured for four hours in the presence, or absence, of wild type human IL-2 (circles) or human “Switch” IL-2 (triangles) at pH7.4 (red and solid lines) or pH6.4 (lime and dashed lines). Blots of one representative experiment are shown. Line graphs shows the mean ± SD of two biological replicates. **d,** Impact of the IL-2 “Switch” cytokine on expansion and viability of human CD8^+^ T cells. Human CD8^+^ T cells were cultured for five days in the presence, or absence, of wild type human IL-2 (circles) or human “Switch” IL-2 (triangles) at pH7.4 (red and solid lines) or pH6.4 (lime and dashed lines). Line graphs shows the mean ± SD of two biological replicates. **e,** CD25 expression levels upon knock out using CRISPR/Cas9. OT-I x CRISPR/Cas9 CTLs were transduced with retroviruses expressing a negative control sgRNA or a CD25 sgRNA. Upon expansion and Thy1.1 enrichment, CD25 expression was analyzed by flow cytometry. One representative histogram is shown. Bar graph shows the percentage of CD25^+^ CTLs ± SEM of three biological replicates out of two independent experiments. **f,** Impact of pH and CD25 on IL-2/IL-2R binding. IL-2/IL-2R binding was determined by incubating cells used in (**e**) with biotinylated IL-2 at various pH. Line graph on the left shows IL-2/IL-2R binding normalized to the condition sgCTRL pH7.4 ± SEM of three biological replicates out of two independent experiments. Line graph on the right displays the same results normalized to the respective sgRNA at pH7.4. **g,** Impact of pH and CD25 on IL-2R signaling. The same cells as used in (**e,f**) were cultured for four hours in the presence, or absence, of murine IL-2 at pH7.4 or pH6.6. Bar graphs shows the mean + SEM of three biological replicates from two independent experiments. **h,** Impact of the IL-2 “Switch” cytokine on expansion and viability. The same cells as used in (**e,f**) were cultured for four days in the presence, or absence, of murine IL-2 at pH7.4 or pH6.6. Bar graphs shows the mean + SEM of three biological replicates from two independent experiments.

**Extended Data Fig. 7:**
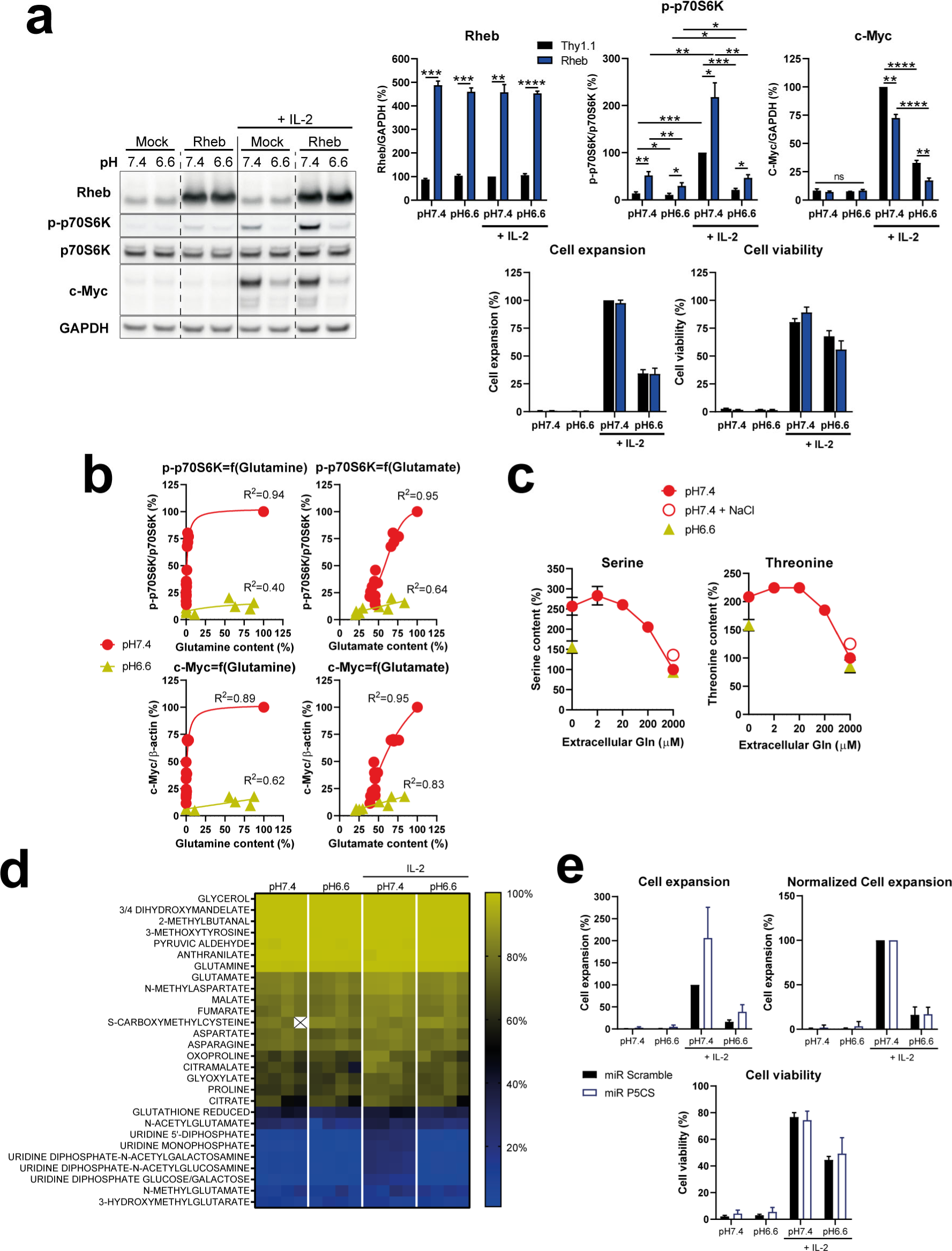
Metabolic incorporation of glutamine, and impact of P5CS-mediated metabolite conversion on CTL proliferation and viability. **a,** Rheb overexpression improves mTORC1 activity. OT-I CTLs were transduced with a control (Mock) or a Rheb encoding retrovirus, and were cultured for 4 hours (western blot analyses) or 4 days (cell expansion and viability) at pH7.4 or pH6.6 with, or without, exogenous murine IL-2 (200 IU/mL). One representative western blot is shown. Bar graphs show the mean phosphorylation status, or total levels, of the indicated molecule relative to the total protein of interest, or to GAPDH, normalized to the condition “pH7.4 + IL-2 Mock” +SEM of four biological replicates from two independent experiments. ns: not significant, *p<0.05, **p<0.01, ***p<0.001, ****p<0.0001 (Student’s paired *t*-test). **b,** mTORC1/c-Myc as a function of intracellular levels of glutamine/glutamate. OT-I CTLs were cultured for 4 hours with exogenous murine IL-2 (200 IU/mL) at pH7.4 (red circles) or pH6.6 (lime triangles) in the presence of various quantities of glutamine. Results show the individual values of p-p70S6K or c-Myc as a function of intracellular amino acid content normalized to the condition “pH7.4 + 2000μM Gln” from four biological replicates out of two independent experiments. A correlation curve and R^2^ for each pH is displayed. **c,** Dose-dependent impact of exogenous glutamine on intracellular serine and threonine levels. OT-I CTLs were cultured for 4 hours with exogenous murine IL-2 (200 IU/mL) at pH7.4 (red circles) or pH6.6 (lime triangles) in the presence of various quantities of glutamine, and in presence or absence of NaCl 31mM. Results show the mean intracellular amino acid content normalized to the condition “pH7.4 + 2000μM Gln” ± SEM of four biological replicates out of two independent experiments. **d,** Metabolites incorporating extracellular glutamine. OT-I CTLs were cultured for four hours with isotopic ^13^C-glutamine (2mM) in the presence, or absence, of exogenous murine IL-2 (200 IU/mL) at pH7.4 or pH6.6. Heat map shows the proportion of the indicated intracellular amino acid that incorporated ^13^C using a color code, as outlined on the ladder. Each condition has four biological replicates from two independent experiments. Empty square with a cross inside means the metabolite was not detected. Metabolites are ordered based on their mean at pH7.4 in the presence of IL-2, and those with less than 10% of ^13^C incorporation are not shown. **e,** P5CS knockdown does not restore CTL proliferation and viability at low pH. OT-I CTLs transduced with a control miR (Scramble) or a P5CS-targeting miR were cultured for 3 days with, or without, exogenous murine IL-2 (200 IU/mL) at pH7.4 or pH6.6. Bar graph on the top and left hand displays mean CTL expansion normalized to the condition “pH7.4 + IL-2” + SEM of four biological replicates from two independent experiments. Bar graph on the top and right hand displays mean CTL expansion normalized to the condition “pH7.4 + IL-2” per corresponding miR + SEM of four biological replicates from two independent experiments. Bar graph on the bottom shows mean CTL viability + SEM of four biological replicates from two independent experiments.

**Extended Data Fig. 8:**
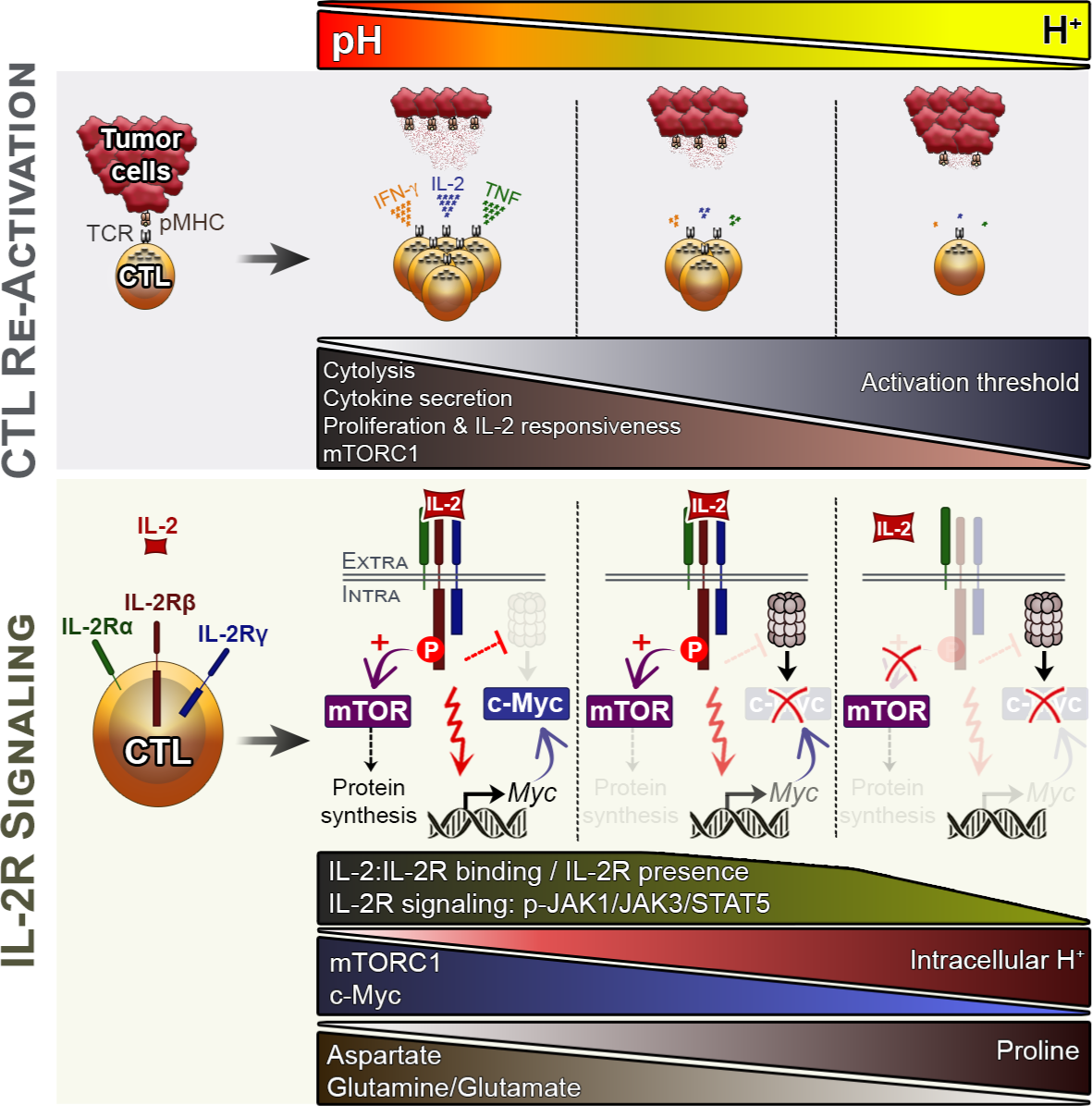
Graphical conclusion.

## References

1. Dudley, M.E. & Rosenberg, S.A. Adoptive-cell-transfer therapy for the treatment of patients with cancer. Nat Rev Cancer 3, 666–675 (2003).

2. Zhang, L. et al. Intratumoral T cells, recurrence, and survival in epithelial ovarian cancer. N Engl J Med 348, 203–213 (2003).

3. Wang, M., Yin, B., Wang, H.Y. & Wang, R.F. Current advances in T-cell-based cancer immunotherapy. Immunotherapy 6, 1265–1278 (2014).

4. Durgeau, A., Virk, Y., Corgnac, S. & Mami-Chouaib, F. Recent Advances in Targeting CD8 T-Cell Immunity for More Effective Cancer Immunotherapy. Front Immunol 9, 14 (2018).

5. Anderson, K.G., Stromnes, I.M. & Greenberg, P.D. Obstacles Posed by the Tumor Microenvironment to T cell Activity: A Case for Synergistic Therapies. Cancer Cell 31, 311–325 (2017).

6. Darvin, P., Toor, S.M., Sasidharan Nair, V. & Elkord, E. Immune checkpoint inhibitors: recent progress and potential biomarkers. Exp Mol Med 50, 1–11 (2018).

7. Vigano, S. et al. Targeting Adenosine in Cancer Immunotherapy to Enhance T-Cell Function. Front Immunol 10, 925 (2019).

8. Chang, C.H. et al. Metabolic Competition in the Tumor Microenvironment Is a Driver of Cancer Progression. Cell 162, 1229–1241 (2015).

9. Salmon, H. et al. Matrix architecture defines the preferential localization and migration of T cells into the stroma of human lung tumors. J Clin Invest 122, 899–910 (2012).

10. Vuillefroy de Silly, R., Dietrich, P.Y. & Walker, P.R. Hypoxia and antitumor CD8(+) T cells: An incompatible alliance? Oncoimmunology 5, e1232236 (2016).

11. Huber, V. et al. Cancer acidity: An ultimate frontier of tumor immune escape and a novel target of immunomodulation. Semin Cancer Biol 43, 74–89 (2017).

12. Warburg, O. On the origin of cancer cells. Science 123, 309–314 (1956).

13. Tannock, I.F. & Rotin, D. Acid pH in tumors and its potential for therapeutic exploitation. Cancer Res 49, 4373–4384 (1989).

14. Swietach, P., Vaughan-Jones, R.D. & Harris, A.L. Regulation of tumor pH and the role of carbonic anhydrase 9. Cancer Metastasis Rev 26, 299–310 (2007).

15. Wu, H. et al. T-cells produce acidic niches in lymph nodes to suppress their own effector functions. Nat Commun 11, 4113 (2020).

16. Taylor, A.C. Responses of cells to pH changes in the medium. J Cell Biol 15, 201–209 (1962).

17. Lardner, A. The effects of extracellular pH on immune function. J Leukoc Biol 69, 522–530 (2001).

18. Irving, M., Vuillefroy de Silly, R., Scholten, K., Dilek, N. & Coukos, G. Engineering Chimeric Antigen Receptor T-Cells for Racing in Solid Tumors: Don’t Forget the Fuel. Front Immunol 8, 267 (2017).

19. Calcinotto, A. et al. Modulation of microenvironment acidity reverses anergy in human and murine tumor-infiltrating T lymphocytes. Cancer Res 72, 2746–2756 (2012).

20. Pilon-Thomas, S. et al. Neutralization of Tumor Acidity Improves Antitumor Responses to Immunotherapy. Cancer Res 76, 1381–1390 (2016).

21. Wang, X., Rickert, M. & Garcia, K.C. Structure of the quaternary complex of interleukin-2 with its alpha, beta, and gammac receptors. Science 310, 1159–1163 (2005).

22. Liao, W., Lin, J.X. & Leonard, W.J. Interleukin-2 at the crossroads of effector responses, tolerance, and immunotherapy. Immunity 38, 13–25 (2013).

23. Ross, S.H. & Cantrell, D.A. Signaling and Function of Interleukin-2 in T Lymphocytes. Annu Rev Immunol 36, 411–433 (2018).

24. Laplante, M. & Sabatini, D.M. mTOR signaling at a glance. J Cell Sci 122, 3589–3594 (2009).

25. Saxton, R.A. & Sabatini, D.M. mTOR Signaling in Growth, Metabolism, and Disease. Cell 168, 960–976 (2017).

26. Aramburu, J., Ortells, M.C., Tejedor, S., Buxade, M. & Lopez-Rodriguez, C. Transcriptional regulation of the stress response by mTOR. Sci Signal 7, re2 (2014).

27. Nikonorova, I.A. et al. Time-resolved analysis of amino acid stress identifies eIF2 phosphorylation as necessary to inhibit mTORC1 activity in liver. J Biol Chem 293, 5005–5015 (2018).

28. Xie, J. et al. cAMP inhibits mammalian target of rapamycin complex-1 and -2 (mTORC1 and 2) by promoting complex dissociation and inhibiting mTOR kinase activity. Cell Signal 23, 1927–1935 (2011).

29. Gaggero, S., et al. IL-2 is inactivated by the acidic pH environment of tumors enabling engineering of a pH-selective mutein. Sci Immunol 7, eade5686 (2022).

30. Farrell, A.S. & Sears, R.C. MYC degradation. Cold Spring Harb Perspect Med 4 (2014).

31. Garami, A. et al. Insulin activation of Rheb, a mediator of mTOR/S6K/4E-BP signaling, is inhibited by TSC1 and 2. Mol Cell 11, 1457–1466 (2003).

32. Angarola, B. & Ferguson, S.M. Coordination of Rheb lysosomal membrane interactions with mTORC1 activation. F1000Res 9 (2020).

33. Koivusalo, M. et al. Amiloride inhibits macropinocytosis by lowering submembranous pH and preventing Rac1 and Cdc42 signaling. J Cell Biol 188, 547–563 (2010).

34. Rogala, K.B. et al. Structural basis for the docking of mTORC1 on the lysosomal surface. Science 366, 468–475 (2019).

35. Buck, M.D., Sowell, R.T., Kaech, S.M. & Pearce, E.L. Metabolic Instruction of Immunity. Cell 169, 570–586 (2017).

36. Wolfson, R.L. & Sabatini, D.M. The Dawn of the Age of Amino Acid Sensors for the mTORC1 Pathway. Cell Metab 26, 301–309 (2017).

37. Loftus, R.M. et al. Amino acid-dependent cMyc expression is essential for NK cell metabolic and functional responses in mice. Nat Commun 9, 2341 (2018).

38. Jewell, J.L. et al. Metabolism. Differential regulation of mTORC1 by leucine and glutamine. Science 347, 194–198 (2015).

39. Fuchs, B.C. & Bode, B.P. Amino acid transporters ASCT2 and LAT1 in cancer: partners in crime? Semin Cancer Biol 15, 254–266 (2005).

40. Scalise, M., Pochini, L., Console, L., Losso, M.A. & Indiveri, C. The Human SLC1A5 (ASCT2) Amino Acid Transporter: From Function to Structure and Role in Cell Biology. Front Cell Dev Biol 6, 96 (2018).

41. Altman, B.J., Stine, Z.E. & Dang, C.V. From Krebs to clinic: glutamine metabolism to cancer therapy. Nat Rev Cancer 16, 619–634 (2016).

42. Yoo, H.C., Yu, Y.C., Sung, Y. & Han, J.M. Glutamine reliance in cell metabolism. Exp Mol Med 52, 1496–1516 (2020).

43. Cormerais, Y. et al. The glutamine transporter ASCT2 (SLC1A5) promotes tumor growth independently of the amino acid transporter LAT1 (SLC7A5). J Biol Chem 293, 2877–2887 (2018).

44. Fischer, K. et al. Inhibitory effect of tumor cell-derived lactic acid on human T cells. Blood 109, 3812–3819 (2007).

45. Nakagawa, Y. et al. Effects of extracellular pH and hypoxia on the function and development of antigen-specific cytotoxic T lymphocytes. Immunol Lett 167, 72–86 (2015).

46. Bosticardo, M. et al. Biased activation of human T lymphocytes due to low extracellular pH is antagonized by B7/CD28 costimulation. Eur J Immunol 31, 2829–2838 (2001).

47. Mendler, A.N. et al. Tumor lactic acidosis suppresses CTL function by inhibition of p38 and JNK/c-Jun activation. Int J Cancer 131, 633–640 (2012).

48. Brand, A. et al. LDHA-Associated Lactic Acid Production Blunts Tumor Immunosurveillance by T and NK Cells. Cell Metab 24, 657–671 (2016).

49. Pollizzi, K.N. & Powell, J.D. Regulation of T cells by mTOR: the known knowns and the known unknowns. Trends Immunol 36, 13–20 (2015).

50. Damaghi, M., Wojtkowiak, J.W. & Gillies, R.J. pH sensing and regulation in cancer. Front Physiol 4, 370 (2013).

51. Jewell, J.L. et al. GPCR signaling inhibits mTORC1 via PKA phosphorylation of Raptor. Elife 8 (2019).

52. Balgi, A.D. et al. Regulation of mTORC1 signaling by pH. PLoS One 6, e21549 (2011).

53. Faes, S. et al. Acidic tumor microenvironment abrogates the efficacy of mTORC1 inhibitors. Mol Cancer 15, 78 (2016).

54. Walton, Z.E. et al. Acid Suspends the Circadian Clock in Hypoxia through Inhibition of mTOR. Cell 174, 72–87 e32 (2018).

55. Wang, R. et al. The transcription factor Myc controls metabolic reprogramming upon T lymphocyte activation. Immunity 35, 871–882 (2011).

56. Gnanaprakasam, J.N. & Wang, R. MYC in Regulating Immunity: Metabolism and Beyond. Genes (Basel) 8 (2017).

57. Marchingo, J.M., Sinclair, L.V., Howden, A.J. & Cantrell, D.A. Quantitative analysis of how Myc controls T cell proteomes and metabolic pathways during T cell activation. Elife 9 (2020).

58. Newsholme, E.A., Crabtree, B. & Ardawi, M.S. Glutamine metabolism in lymphocytes: its biochemical, physiological and clinical importance. Q J Exp Physiol 70, 473–489 (1985).

59. DeBerardinis, R.J. et al. Beyond aerobic glycolysis: transformed cells can engage in glutamine metabolism that exceeds the requirement for protein and nucleotide synthesis. Proc Natl Acad Sci U S A 104, 19345–19350 (2007).

60. Hammami, I., Chen, J., Bronte, V., DeCrescenzo, G. & Jolicoeur, M. L-glutamine is a key parameter in the immunosuppression phenomenon. Biochem Biophys Res Commun 425, 724–729 (2012).

61. Hayat, S. et al. Role of proline under changing environments: a review. Plant Signal Behav 7, 1456–1466 (2012).

62. Szabados, L. & Savoure, A. Proline: a multifunctional amino acid. Trends Plant Sci 15, 89–97 (2010).

63. Coren, L.V., Jain, S., Trivett, M.T., Ohlen, C. & Ott, D.E. Production of retroviral constructs for effective transfer and expression of T-cell receptor genes using Golden Gate cloning. Biotechniques 58, 135–139 (2015).

64. Chang, K., Elledge, S.J. & Hannon, G.J. Lessons from Nature: microRNA-based shRNA libraries. Nat Methods 3, 707–714 (2006).

65. Pelossof, R. et al. Prediction of potent shRNAs with a sequential classification algorithm. Nat Biotechnol 35, 350–353 (2017).

66. Labun, K., Montague, T.G., Gagnon, J.A., Thyme, S.B. & Valen, E. CHOPCHOP v2: a web tool for the next generation of CRISPR genome engineering. Nucleic Acids Res 44, W272–276 (2016).

67. Chen, B. et al. Dynamic imaging of genomic loci in living human cells by an optimized CRISPR/Cas system. Cell 155, 1479–1491 (2013).

68. Naviaux, R.K., Costanzi, E., Haas, M. & Verma, I.M. The pCL vector system: rapid production of helper-free, high-titer, recombinant retroviruses. J Virol 70, 5701–5705 (1996).

69. Jordan, M., Schallhorn, A. & Wurm, F.M. Transfecting mammalian cells: optimization of critical parameters affecting calcium-phosphate precipitate formation. Nucleic Acids Res 24, 596–601 (1996).

70. De Jong, M.O., Rozemuller, H., Bauman, J.G. & Visser, J.W. Biotinylation of interleukin-2 (IL-2) for flow cytometric analysis of IL-2 receptor expression. Comparison of different methods. J Immunol Methods 184, 101–112 (1995).

71. Gallart-Ayala, H. et al. A global HILIC-MS approach to measure polar human cerebrospinal fluid metabolome: Exploring gender-associated variation in a cohort of elderly cognitively healthy subjects. Anal Chim Acta 1037, 327–337 (2018).

72. Midani, F.S., Wynn, M.L. & Schnell, S. The importance of accurately correcting for the natural abundance of stable isotopes. Anal Biochem 520, 27–43 (2017).

73. Roci, I. et al. Metabolite Profiling and Stable Isotope Tracing in Sorted Subpopulations of Mammalian Cells. Anal Chem 88, 2707–2713 (2016).

74. Teav, T. et al. Merged Targeted Quantification and Untargeted Profiling for Comprehensive Assessment of Acylcarnitine and Amino Acid Metabolism. Anal Chem 91, 11757–11769 (2019).

